# Towards measurements of absolute membrane potential in Bacillus subtilis using fluorescence lifetime

**DOI:** 10.1101/2024.06.13.598880

**Authors:** Debjit Roy, Xavier Michalet, Evan W. Miller, Kiran Bharadwaj, Shimon Weiss

**Affiliations:** UCLA-DOE Institute for Genomics and Proteomics, Department of Biological Chemistry, University of California at Los Angeles, Los Angeles, CA 90095, USA; Department of Chemistry and Biochemistry, University of California at Los Angeles, Los Angeles, CA 90095, USA; Departments of Chemistry, Molecular and Cell Biology, Helen Wills Neuroscience Institute, University of California at Berkeley, CA 94720, USA; Department of Physiology, University of California at Los Angeles, Los Angeles, CA 90095, USA; California Nano Systems Institute, University of California at Los Angeles, Los Angeles, CA 90095, USA; Department of Physics, Institute for Nanotechnology and Advanced Materials, Bar-Ilan University, Ramat-Gan 52900, Israel

**Keywords:** Bacterial Bioelectricity, Membrane Potential, Optical Electrophysiology, VoltageFluor, Fluorescence Lifetime Imaging Microscopy, Phasor Analysis

## Abstract

Membrane potential (MP) changes can provide a simple readout of bacterial functional and metabolic state or stress levels. While several optical methods exist for measuring fast changes in MP in excitable cells, there is a dearth of such methods for absolute and precise measurements of steady-state membrane potentials (MPs) in bacterial cells. Conventional electrode-based methods for the measurement of MP are not suitable for calibrating optical methods in small bacterial cells. While optical measurement based on Nernstian indicators have been successfully used, they do not provide absolute or precise quantification of MP or its changes. We present a novel, calibrated MP recording approach to address this gap. In this study, we used a fluorescence lifetime-based approach to obtain a single-cell resolved distribution of the membrane potential and its changes upon extracellular chemical perturbation in a population of bacterial cells for the first time. Our method is based on (i) a unique VoltageFluor (VF) optical transducer, whose fluorescence lifetime varies as a function of MP via photoinduced electron transfer (PeT) and (ii) a quantitative phasor-FLIM analysis for high-throughput readout. This method allows MP changes to be easily visualized, recorded and quantified. By artificially modulating potassium concentration gradients across the membrane using an ionophore, we have obtained a *Bacillus subtili*s-specific MP versus VF lifetime calibration and estimated the MP for unperturbed *B. subtilis* cells to be −65 mV (in MSgg), −127 mV (in M9) and that for chemically depolarized cells as −14 mV (in MSgg). We observed a population level MP heterogeneity of ∼6-10 mV indicating a considerable degree of diversity of physiological and metabolic states among individual cells. Our work paves the way for deeper insights into bacterial electrophysiology and bioelectricity research.

## INTRODUCTION

Cellular communication, pattern formation and behavior are governed not only by chemical cues (e.g., chemo-attractants, ion gradients, protein and RNA levels), but also by membrane electrical potentials (1). The term bioelectricity is used to describe phenomena including changes in membrane potential (MP) that mediate and regulate electrical signaling between neighboring cells in a tissue or in a biofilm (2). The membrane potential is defined as the electric potential difference across the membrane lipid bilayer and is maintained by the careful balance between the diffusion pressure created by ionic concentration gradient across the membrane and electrical forces arising from the separation of charges and the selective permeability of the membrane to certain ions. Additionally, charged/zwitterionic lipids, lipid-water dipole alignment, and fixed intracellular charges (DNA, RNA, ribosomes, and proteins) also contribute to the MP (2).

The MP is an intrinsic characteristic of each cell type (3) which the cell tries to maintain to achieve homeostasis. Changes in MP is a mere reflection of changes in the cell’s functional state, often trigger reflex responses due to external stimuli or follow downstream signaling pathways in various physiological processes such as cellular proliferation and differentiation (4). MP changes drive various essential bacterial activity such as energy generation (5), nutrient uptake and transport (6), cell division(7), metabolism (8), osmoregulation (9), mechano-sensation (10), motility (11) and antibiotic resistance (12). Intercellular communication and co-ordination between microbial cells during metabolism and stress response are accompanied, and possibly mediated by, dynamic MP changes followed by ion translocation (electro-chemical communication). Recent studies have shown that MP changes mediate cell-cell signaling in *Bacillus subtilis* (*B. subtilis*) bacterial communities (13), co-ordinate metabolic oscillations radially within biofilms, broadcasting metabolic status amongst cells located within different parts of biofilms (14), and enable dormant spores to integrate environmental signals (15). In summary, the role of membrane potential in all aspects of bacterial physiology cannot be understated. Understanding the relationship of membrane potential changes with cellular physiological processes is crucial. These insights could aid in the development of diagnostic tools for detecting bacterial infections and assessing the efficacy of antibiotic treatments. They could also help in creating solutions for combating antibiotic resistance and designing bacterial strains for industrial applications, such as biofuel production and bioremediation. This underscores the need for robust and accurate methods to quantify membrane potential.

### Optical Estimation of Resting Membrane Potential: problem and potential solution

However, calibrated measurements of absolute MPs and their changes are challenging. Past electrophysiological studies have mainly focused on specialized excitable mammalian cells (neurons, cardiomyocytes, muscle cells, and some secretory cells) and has led to the development of a fundamental framework and models for understanding bio-electricity. Slow changes in MP are common to most tissues and bacterial cells, in contrast to action potentials (APs), *i*.*e*. fast MP changes, which are limited to specialized excitable cells. However, tools to *quantitatively* measure slow changes in MP trail behind tools used to simply detect Aps (16-19). In mammalian cells, electrode-based methods (patch-clamp or micro-electrode arrays) are considered the gold-standard for precise measurements of APs and MPs (16-20). These techniques are sensitive, provide sub-ms temporal resolution, but have limited throughput and spatial resolution. Direct electrical potential measurements using microelectrodes has been achieved on artificially enlarged (genetically /pharmacologically) *E. coli* cells (21). However, electrode-based tools are not suitably implementable for small sized bacterial cells. Alternatively, bacterial MP measurements resorted on the use of small molecule Nernstian indicators that exhibits MP change induced reversible accumulation-redistribution and monitored by radioactivity (TPMP^+^, TPP^+^) (22) or fluorescence (ThT, TMRM, Rh123) measurements (23). Fluorescent Nernstian indicators have been until recently the main tool for studying bacterial membrane potential, as they allow live recording and long-term studies of single cells or biofilms. These are in general cationic, membrane-permeable dyes, with a low membrane binding affinity. The equilibrium concentrations of such an indicator on both sides of the membrane are related to the value of the MP (noted *V*_*m*_) by the Nernst equation:

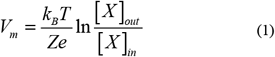

where *Ze* is the electric charge of indicator X, *k*_*B*_ is the Boltzmann constant, *T* the temperature and [*X*]_*out/in*_ the outer/inner indicator concentrations, respectively. Assuming a linear dependence of fluorescence intensity with concentration, MP changes therefore translate into measurable logarithmic changes in the ratio of extracellular to cytoplasmic radioactivity or fluorescence intensities.

Fluorescent Nernstian indicators have been employed to measure bacterial MP, providing insight into electrical spiking in *E. coli* (24), metabolic oscillations in *B. subtilis* biofilms (13), external electrical stimuli induced bacterial MP fluctuations (25), and values of resting MPs ranging from −140 mV to −75 mV (21,26-28).

However, problems exist for Nernstian indicators. At high indicator concentration (>100 µM), cellular MP is likely perturbed (23), and even low concentrations (>10 μM) of dye substantially slow the elongation rate of *B. subtilis* and *E. coli* (29). Another approach to MP measurement in bacteria is the use of genetically engineered voltage indicators (GEVIs) such as PROPS (24,30,31). This powerful technique requires modifying the cell’s genome, making it inappropriate for wild type cell studies, and using an inducer to control expression of the fluorescent GEVI, adding to the uncertainty and complexity of the measurements.

Like all indirect measurements, optical (or radiolabel-based) recordings of MP changes require additional calibration for absolute quantification. However, these measurements are often subjected to confounding effects due to variations in probe concentration, non-uniform microenvironment, probe loading efficiency, variable illumination intensity or background, and photobleaching. Careful control experiments are needed to minimize artifacts due to these variations. In particular, it is very hard to measure small MP changes using Nernstian indicators. For these reasons, better tools are desirable for studying bacterial bioelectricity in real-time, with appropriate spatial resolutions, and voltage sensitivity.

To address these limitations, we explored the use of VoltageFluor dye VF2.1.Cl, a lipophilic dye characterized by a membrane potential-dependent fluorescence lifetime. Fluorescence lifetime imaging microscopy (FLIM) is a powerful tool to monitor the nanoscale environment of biomolecules. The fluorescence lifetime *τ* is an intrinsic molecular property of a fluorophore and is only sensitive to alterations in intramolecular structure (conformational changes) and intermolecular interaction with its local electronic microenvironment. The measured fluorescence lifetime (but not its precision) is independent from the recorded fluorescence intensity and to a large extent from fluorophore concentration, simplifying calibration and reproducibility (32). *τ* is an excellent indicator for local changes in temperature, polarity, viscosity, pH, ionic strength etc. Spatiotemporal mapping of these properties could be acquired via FLIM (33-35). Quantification of MPs by measurement of the fluorescence lifetime of voltage sensitive dyes (VSDs) is therefore an attractive alternative to current approaches.

VoltageFluors have been extensively characterized in eukaryotic cells (36-40). Here, we report the first use of VoltageFluors in bacterial cells. We used *B. subtilis*, a rodshaped, Gram-positive model bacterium (i.e., with a single membrane), and VoltageFluor (VF) dyes as a proof-of-concept testbed. Because patch-clamp is not an option in bacteria, we controlled the MP by addition of chemicals extracellularly and limited ourselves to equilibrium measurements of VF’s fluorescence lifetime after a fixed incubation time. Our results are organized as follows. We first report the photophysical characterization of VF dyes in solution and in live *B. subtilis* membrane. We discuss the effect of MP-induced changes on the photoinduced electron transfer (PeT) kinetic rate affecting the fluorescence lifetime of the VF indicators using a simplified model introduced by Li (41). We then briefly describe the phasor analysis of fluorescence lifetime used in our work and compare it with the conventional non-linear least-square (weighted) fitting (NLSF) approach. Next, we describe the method used to obtain a calibrated MP versus lifetime relation in *B. subtilis*. We report average VF2.1Cl fluorescence lifetime changes observed at the single-cell level and *B. subtilis*’s MP estimated in unperturbed and depolarizing chemical conditions. We further probe the effect of medium osmolality on MP. Color-coded fluorescence lifetime maps visualizing MP changes are discussed in the next to last section, before a summary and critical discussion of our results are presented in the last section. Materials and methods are briefly described after the main text.

### Photophysical characterization of VoltageFluor lifetime in solution and in the membrane of *B. subtilis*

In this work, we used two dyes to measure bacterial MP (Figure 1a): (i) VF2.1.Cl, a voltage-sensitive fluorophore (36,37,42), and (ii) VF2.0.Cl, an isostructural control dye which is not voltage sensitive. Both VF2.1.Cl (voltage-sensitive) and VF2.0.Cl (insensitive) localize to the plasma membrane of mammalian cells. VF2.1.Cl displays voltage- and electric field-dependent fluorescence (43) arising from a photoinduced electron transfer (PeT) from the electron rich aniline into the excited state of the fluorescein fluorophore (Figure 1a) (36,37,42). At hyperpolarized potentials (negative inside), the rate of PeT is accelerated by alignment with the electric field of the membrane, making the dye dim (Figure 1b). At depolarized potentials (positive inside), the rate of PeT is slowed by opposing electric field, making the dye bright (Figure 1c). VF2.0.Cl shows no voltage-dependent fluorescence, in either intensity or lifetime modes (38,44).

**FIGURE 1.**
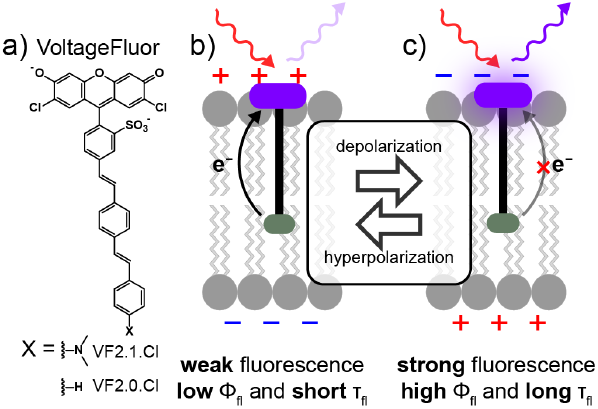
Voltage-sensitive fluorophore VF2.1.Cl. VF2.1.Cl senses MP via photoinduced-electron transfer. In mammalian cells, in the −80 mV to +80 mV range of transmembrane voltage, it exhibits a relative fluorescence intensity (ΔF/F) change of ∼27% per 100 mV and Δt_avg_ ∼3 ps/mV. a) When the membrane gets hyperpolarized, PeT is increased and fluorescence is quenched, b) When the membrane gets depolarized, PeT is reduced and fluorescence enhanced c) Chemical molecular structure of VF2.1.Cl (adapted from Ref. 40).

The mechanism of voltage sensing relies on dependence of the sensor’s intrinsic PeT process on the local electric field *E*. PeT is a non-radiative relaxation pathway from the dye’s first excited state S_1_ to a charge-separated state S_2_, which competes with radiative decay (characterized by fluorescence rate *k*_*f*_) and other non-radiative processes (rate *k*_*nr*_), as discussed in Supplementary Note 6 and illustrated in the Jablonski diagram shown in Supplementary Figure 15. In the absence of PeT, a dye’s fluorescence is normally characterized by a mono-exponential relaxation down to the ground state S_0_ with rate *k*_*10*_ = *k*_*f*_ *+ k*_*nr*_ equal to the inverse of the measured fluorescence lifetime. This behavior was observed by the Miller group for VF2.0.Cl, which did not show voltage-dependent fluorescence intensity, in mammalian cells (38,44).

In the presence of PeT, hyperpolarized membranes (membranes with more negative membrane potential difference than the resting membrane potential) resulted in enhanced PeT, which led to increased nonradiative relaxation from the dye’s excited state S_1_ to the charge-separated state S_2_ (rate *k*_*12*_ = *k*_*PeT*_), followed by relaxation to the ground state S_0_ (with rate *k*_*20*_), resulting in an overall shorter average lifetime and dimmer fluorescence (Figure 1a). Inversely, depolarized membranes resulted in reduced PeT and longer lifetime, *i*.*e*. brighter emission (Figure 1b) (38,39,44-47).

Using the 3-state model described in Supplementary Note 6, one obtains the following result for the amplitude-averaged lifetime ⟨*τ*⟩_*a*_ (Supplementary Eq. S11) and quantum yield *Y*_*f*_ (Supplementary Eq. S10):

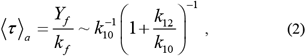

where the membrane potential (*V*_*m*_) dependence of ⟨ *τ* ⟩_*a*_ is encompassed in the electric field *E*-dependence of *k*_*12*_ via *V*_*m*_ *= E/d*, where *d* is the membrane thickness (Supplementary Eqs. S7-S8). This dependence can be used to measure the membrane potential, after calibration of the VF’s response to a series of externally applied membrane potentials (38,48). The voltage sensitivity of various VF dyes has been previously characterized in cultured cells, dissociated mammalian neurons, and *ex vivo* leech ganglia under patchclamp, which allowed precise control of the membrane potential (37,38,44). Woodford *et al*. observed various types of dependence of the relative intensity change Δ*F/F =* Δ*Y/Y* on *V*_*m*_ for a variety of VF dyes. In particular, VF2.1.Cl exhibited a linear dependence of Δ*F/F* on *V*_*m*_ in the ±80 mV range in HEK cells incubated in HBSS buffer.

These characteristics made VF2.1.Cl an ideal VF dye to test for bacterial membrane potential measurements. We chose *B. subtilis* as a model Gram-positive (single membrane) bacterial species, as it has been extensively characterized in the literature and is non-pathological.

We first compared the fluorescence lifetime of both VF2.1.Cl and VF2.0.Cl in solution and in bacterial membranes. In solution, VF2.1.Cl exhibited a faster fluorescence decay than VF2.0.Cl. due to the existence of an intrinsic PeT channel, absent in VF2.0.Cl (Supplementary Figure 1). Specifically, VF2.0.Cl exhibited a single-exponential decay (<τ>_2.0,DMSO_ = 3.21 ns), whereas VF2.1.Cl was characterized by a clear bi-exponential decay (τ_1_ = 2.90 ns, τ_2_ = 0.81 ns; <τ>_2.1,DMSO_ = 1.53 ns) as obtained by NLSF analysis (Supplementary Table 2).

The membrane of *B. subtilis* was readily stained with both dyes (see Supplementary Note 1 for bacteria culture, growth and membrane staining protocols). The fluorescence signal appears limited to the membrane based on a comparison of the observed intensity profile to a simple model accounting for the microscope’s optical characteristics and defocusing (Figure 2a,b and Supplementary Note 7). The intensity-weighted lifetime map (Figure 2c) obtained by phasor analysis, as described in the next section, indicates a fairly uniform lifetime, with some measurable fluctuations compatible to those expected from signal shot noise, as illustrated in Figure 2d and discussed in a later section. The amplitude-averaged fluorescence lifetime of VF2.0.Cl in the membrane of *B. subtilis* was <τ>_2.0,mb_ = 2.72 ns, while that of VF2.1.Cl was <τ>_2.1,mb_ = 0.74 ns (τ_1_ = 1.87 ns, τ_2_ = 0.32 ns) (Supplementary Table 2). Interestingly, both decays exhibited two exponential components, including that of VF2.0.Cl. For VF2.1.Cl, the reduction of the average lifetime is related to the different environments of the VF and the presence of the transmembrane electric field associated with the membrane, as studied next.

**FIGURE 2B.**
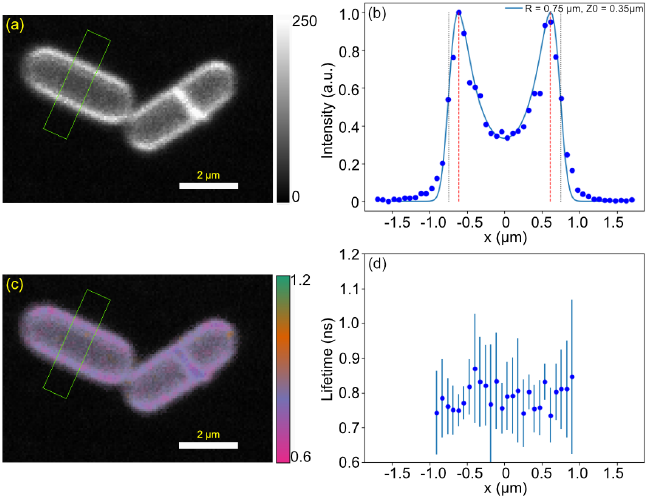
subtilis membrane staining with VF2.1.Cl. (a) Confocal image (72 nm/pixel) of two isolated bacteria. Bar indicates 2 μm. (b) Dots: intensity profile averaged along the short axis of the rectangle ROI shown in a. The blue line corresponds to expected signal from a cylindrical fluorescent object with radius R = 0.75 μm (grey vertical line) and focus z_0_ = 350 nm above the midplane. The apparent radius (red line) is 0.61 μm. (c) Color-coded amplitude-averaged lifetime map. (d) Dots: amplitude-averaged lifetime profile averaged along the short axis of the rectangle ROI shown in a & c. The error bar corresponds to the standard deviation of the lifetime along the short axis of the ROI. The lifetime profile is limited to the pixels with intensity above background.

### Fluorescence lifetime analysis: nonlinear least-square fitting versus phasor approach

Conventionally fluorescence lifetime determinations are performed by TCSPC FLIM followed by nonlinear least-square fitting (NLSF) of the recorded photon arrival time histogram. This is computationally expensive and requires high photon counts/good signal-to-noise ratio (SNR), especially for bi-exponential decays (49,50). We therefore turned to phasor analysis (51-62). Phasor analysis can be as quantitative as NLSF for bi-exponential decays when individual lifetime components are known or can be determined independently (59,62,63). Here, we introduce a heuristic approach (described in: Material & Methods and Supplementary Note 4) to internally define individual single-exponential reference lifetimes from population data. We compared the lifetime of VF2.1.Cl in *B. subtilis* determined by NLSF (741 ± 62 ps, Figure 3a) and by phasor analysis (778 ± 38 ps, Figure 3b). The values are in good agreement, the difference of 37 ps being due to different approximations used in both methods. Based on this agreement, we decided to use the phasor analysis approach in the remainder of this study, due to its speed and insensitivity to signal-to-noise ratio (51-57).

**FIGURE 3.**
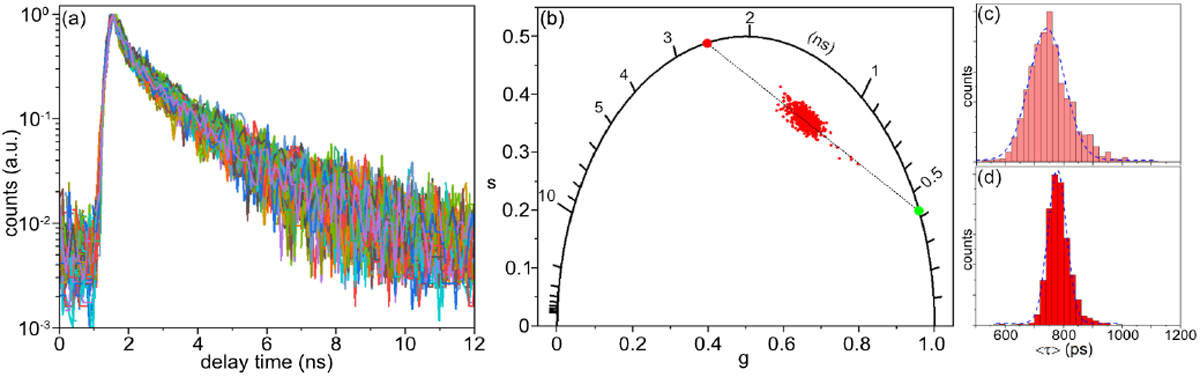
(a) 100 representative single-cell fluorescence decays for a population of *B. subtilis* cells stained with VF2.1.Cl (suspended in MSgg medium). Their fit with two-exponential decay model yields time constants τ_1_ = 1.87 ± 0.16 ns and τ_2_ = 0.32 ± 0.05 ns. (b) Corresponding phasor plot, with the single-exponential references (green and red dots on the UC, τ_1_ = 2.51 ns, τ _2_ = 0.44 ns) and the segment connecting them (dashed line). (c) amplitude-averaged lifetime <τ> histogram obtained via single-cell NLSF analysis (<τ> = 741 ± 62 ps) and (d) amplitude-averaged lifetime <τ> histogram obtained by single-cell phasor analysis (<τ> = 778 ± 32 ps). The two-sample t-test with Welch correction for unequal variance indicates a small but significant difference between the two distributions, which are attributable to small systematic differences resulting from different reference choices in NLSF and phasor analysis.

### Single cell membrane potential calibration of *B. subtilis* via phasor-FLIM measurements

The electric field across the membrane depends on local charge densities on both side of the membrane (whose possible non-homogeneity along the cell periphery is neglected in this work). This local density is connected by a boundary layer to the internal or external bulk ion concentrations, respectively, and can be described by various models, ranging from the simple Gouy-Chapman double-layer model (64) to more sophisticated ones solving the Poisson-Boltzmann equation (e.g. (65-67)). The important result of this type of analysis is that, far away from the membrane, the bulk ion concentrations can be considered constant and, at equilibrium, can be computed using the Donnan model (68) or, when taking the permeabilities *p*_*i*_ of the different ions through the membrane into account, the Goldman-Hodgkin-Katz (GHK) model (2,64):

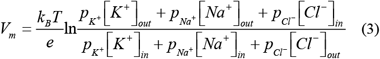

where index *in* (resp. *out*) indicate concentration inside (resp. outside) the cell. Of the 3 monovalent ions (multivalent ions being less permeable are typically neglected) contributing most to the membrane potential (K^+^, Na^+^ and Cl^-^), the first one is the most membrane permeable, leading to a GHK solution which is effectively well approximated by the Nernst potential of K^+^ (Eq. (1)):

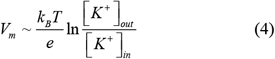

Thus, if [K^+^]_in_ is known for a given external medium composition (with a specific [K^+^]_out_), the membrane potential can be computed using Eq. (4).

However, this estimated membrane potential is not exactly the potential drop across the membrane. There is an electrostatic potential gradient near the membrane due to varying ion concentrations, which reduces the effective membrane potential drop to V_mb_, as discussed in Supplementary Note 8. In most practical cases, the resulting electric field to which the dye is exposed, *E* = *V*_*mb*_/*d* is only slightly smaller (by 1-3%) than that estimated from the Nernst potential *V*_*m*_ (Eq. (4)), justifying neglecting this difference (see Supplementary Note 8 for details). Note that this estimate ignores the possible contribution of charged species residing within the external bacterial wall to the electrostatic potential, a point which will be discussed in the last section.

Patch clamp electrophysiology is impractical for controlling the MP of *B. subtilis*. Therefore, another method was needed to control the MP for a calibration. Since the MP stems from the unequal distribution of ions and charged species across the membrane (mainly K^+^, as discussed previously), one way to modulate the MP is to artificially change ionic concentration gradients across the membrane in conjunction with an ionophore. Ionophores, a class of physiologically active compounds (69-71) that facilitate selective ion transport across cell membranes (bypassing normal ion channels or transporters). They can be used to increase the membrane permeability for specific ions. This, in turn, alters intracellular ionic compositions and ionic concentration gradients across the cell membranes. Here, we used valinomycin, a mobile carrier ionophore, known for selective translocation of K^+^ across biological and synthetic membranes down the electrochemical potential gradient (70,72-77). Valinomycin treatment effectively enhances the K^+^ permeability through the cell membrane, making it impossible for the cell to effectively maintain its resting potential (70,73-77). The accepted mechanism of K^+^ transport by valinomycin involves binding to K^+^ on one side of the membrane, dissolution into the membrane lipid bilayer, followed by transport and release of K^+^ on the other side of the membrane (73-75). The increased permeability of K^+^, coupled with changing [K^+^]_out_, unsettles the existing K^+^ concentration gradient. The cell partially counterbalances the change, but it eventually leads to an altered [K^+^]_in_/[K^+^]_out_ ratio. This new ratio differs from the original one, resulting in a different membrane potential.

We used a previously established *V*_*m*_ *vs* [K^+^]_out_ calibration curve measured in *B. subtilis* using Nernstian radiolabels ([^3^H]triphenylmethylphosphonium bromide, ^3^TPMP^+^) in the presence of valinomycin and varying external concentrations of K^+^ (78). By measuring the inside and outside bulk concentrations of the indicator at equilibrium, [^3^TPMP^+^]_in_ and [^3^TPMP^+^]_out_, Shioi *et al*. were able to correlate [K^+^]_out_ and the membrane potential *V*_*m*_ computed with Eq. (1) for the indicator. Their data, obtained at variable [K^+^]_out_ but constant external osmolality, are reproduced in Figure 4a.

**FIGURE 4:**
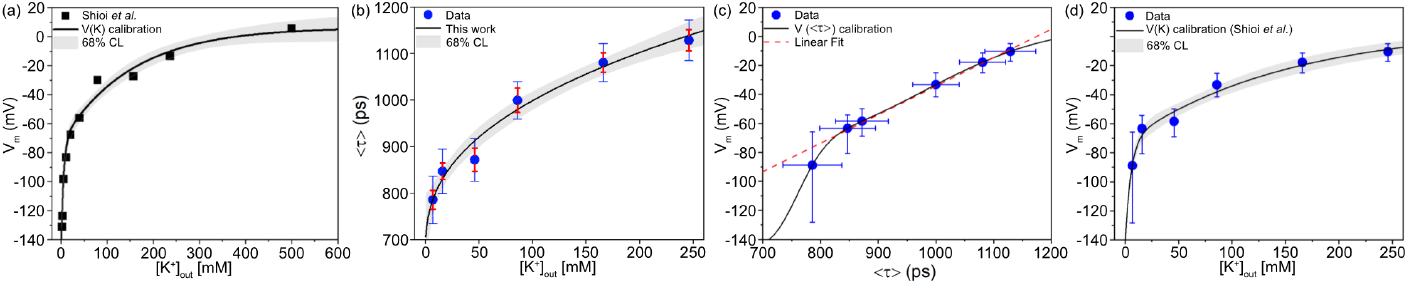
: Calibration of the response of VF2.1.Cl to *B. subtilis* membrane potential changes. (a) *B. subtilis* membrane potential as function of external K^+^ concentration measured with radioactive Nernstian probes in the presence of 10 μM valinomycin (adapted from Figure 3 in Ref. 78). The line is a biexponential fit *V*_*m*_([K^+^]_out_) to the data. (b) Measured averaged fluorescence lifetime <τ> of VF2.1.Cl in the membrane of *B. subtilis* as a function of external K^+^ concentration measured in similar conditions as in (a) (medium osmolality ∼650 mOsm/L). The line is a power law fit with offset <τ>([K^+^]_out_) to the data. The blue vertical error bars indicate the population’s standard deviation of measured fluorescence lifetime and red vertical error bars indicate the estimated shot-noise contribution for the corresponding measurement. (c) *V*_*m*_ *vs* <τ> calibration curve obtained by combining the two fitted relations shown in (a) & (b). The horizontal error bars indicate the standard deviation of the data used for calibration (as shown in b), while the vertical error bars represent the corresponding uncertainty in the membrane potential. The red dashed line corresponds to a linear fit of the data with slope 0.2 mV/ps. (d) *V*_*m*_ values obtained for the [K^+^]_out_ used in the calibration series using the *V*_*m*_([K^+^]_out_) obtained in (a). The error bar was computed as in (c).

We recorded the fluorescence lifetime of VF2.1.Cl at different [K^+^]_out_ in the presence of valinomycin (Figure 4b), exactly reproducing the previous calibration conditions (78). Briefly, *B. subtilis* cells in the mid-exponential phase were treated with 25 µM valinomycin and supplemented with various concentrations of KCl (1 to 240 mM). To maintain constant medium osmolality, NaCl was added to keep the total cation concentration equal to 240 mM (medium osmolality ∼650 mOsm/L). We then constructed a VF2.1.Cl lifetime (<τ>) *vs V*_*m*_ calibration curve (Figure 4c) by combining the previously obtained *V*_*m*_ *vs* [K^+^]_out_ data (78) (Figure 4a) and the measured fluorescence lifetime <τ> *vs* [K^+^]_out_ data (Figure 4b). This *V*_*m*_-<τ> relation is approximately piecewise linear, with a slope of 0.2 mV/ps (5 ps/mV) in the 850 ps – 1.2 ns range, and increasing to up to 0.8 mV/ps (1.2 ps/mV) for lower lifetimes (see Supplementary Note 9 for details).

We turn next to the dispersion of measured lifetime values and its effect on estimated MP. As can be seen in Figure 4d, the calibration relation becomes increasingly uncertain as [K^+^]_out_ decreases. This observation in part reflecting the saturation of VF2.1.Cl’s quenching at very negative membrane potential in B. *subtilis*. However, a significant part of this dispersion can be attributed to the limited SNR of the FLIM data. To assess the contribution of shot noise to the standard deviation of measured lifetime, we have estimated the contribution of shot noise to the phasor-based amplitude-averaged lifetime for groups of cells having similar signal (discussed in Material & Methods and Supplementary Note 5). In Figure 4b, blue vertical error bars represent the total population-level standard deviation of the measured lifetime, while the red error bars represent the population-level standard deviation of the lifetime, if only shot noise contributed to the lifetime dispersion. The results unambiguously show that the population-level distribution of lifetimes is broader than that due to shot noise, suggesting that an additional source of cell-to-cell variation in the measured lifetime is present. Assuming uncorrelated contributions of shot noise and “other sources” (e.g. cell-to-cell physiological, metabolic state variations) to the observed lifetime variance, we estimated the lifetime standard deviation due to these non-shot noise sources, 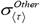, by:

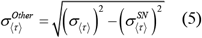

In most measurements, we find that shot noise accounts for a significant part of the observed amplitude-averaged lifetime dispersion (see Supplementary Table 5). For instance, in our calibration series, some measurements show a residual (non-shot noise related) lifetime standard deviation ranging from ∼30 to 50 ps, which corresponds to ∼6-10 mV using the approximate 0.2 mV/ps conversion coefficient of Figure 4c (see Supplementary Table 5). The biological implication of this additional dispersion is the existence of an inherent MP heterogeneity within the bacterial population. Such an intra-population MP variability has been previously observed using Nernstian dyes(23,25) and is also known to exist for other physiological characteristics such as internal pH (79), although this has never been formally characterized. It is likely due to individual cells being in different metabolic or physiological states.

To assess the influence of osmolality (and therefore osmotic pressure) on these results, we performed similar measurements using a different total cation concentration equal to 300 mM (medium osmolality ∼770 mOsm/L instead of 650 mOsm/L). The results, represented in (Figure 5), are globally similar, with a day-to-day variability of about ±100 ps (Figure 5a) equivalent to ±20 mV using the previous calibration (Figure 5b). Day-to-day variability can be attributed to variations in the cells’ physiological state at the time of harvesting prior to media perturbation, staining and imaging and could be improved by a better control of the cell culture’s initial state. When adding increasing concentration of potassium without maintaining a constant osmolality, we observed a comparable increase of lifetime (or membrane depolarization, see Supplementary Figure 2) with increasing external K^+^ concentration. It is worth noting that these differences between measurements across series remain qualitatively similar as extracellular chemical conditions change from the “unperturbed” state to higher external salt concentration. In other words, measurement of a lower lifetime in MSgg in the presence of valinomycin (for instance Series 2 in Figure 5a), will exhibit lower lifetimes throughout the series.

**FIGURE 5:**
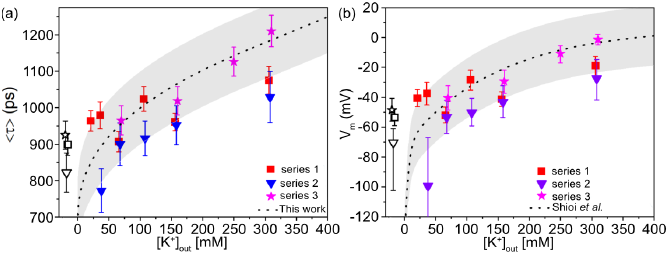
VF2.1.Cl lifetime measurements in *B. subtilis* cells in MSgg media exposed to variable external K^+^ concentration in the presence of 25 μM valinomycin at constant medium osmolality ∼770 mOsm/L. (a) Average lifetime and (b) estimated membrane potential using the calibration curve of Figure 4c. The dashed curves correspond to the curves in Figure 4b and 4d respectively, obtained at 650 mOsm/L external osmolality. The grayed areas correspond to these curves ± 100 ps and ± 20 mV respectively. Black symbols correspond respectively to the measured lifetimes and membrane potentials of *B. subtilis* in MSgg only (no valinomycin or added salts). It is noteworthy that the larger <τ> (resp. *V*_*m*_) is in MSgg only, the larger it is in that series.

Altogether, these experiments confirm that in the presence of valinomycin, *B. subtilis*’ membrane potential is mostly determined by the external potassium concentration, although some variations might be expected depending on the initial cell’s metabolic and physiological state (as discussed earlier).

To elucidate the possible role of potassium concentration, or more generally, of changing external conditions on the fluorescence of VF2.1.Cl itself, we measured its fluorescence decay in MSgg and MSgg with valinomycin and with or without excess KCl, in the absence of cells. While addition of valinomycin (25 μM) only had negligible effect on VF2.1.Cl’s lifetime (<τ>_MSgg,0_ = 2.06 ns *vs* <τ>_MSgg,Val_ = 2.20 ns), addition of excess of KCl (240-300 mM) in presence of valinomycin resulted in a noticeable decrease of VF2.1.Cl’s lifetime (<τ>_MSgg,Val,KCl_ = 1.69 ns, Supplementary Table 3 and Supplementary Figure 3). Specifically, we observed that this variation was mainly due to changes in the respective fractions of the long and short decay components of the decay. The resulting decreased average lifetime (<τ>_MSgg,Val,KCl_ = 1.69 ns) was still much larger than those recorded in the membrane of *B. subtilis* in all our measurements (0.75 ns < <τ>_mb_ < 1.15 ns). This shows that, while VF2.1.Cl’s lifetime in solution is surprisingly sensitive to its ionic environment, once inserted in the membrane of *B. subtilis*, the observed lifetime variations of VF2.1.Cl report on membrane potential changes (see Supplementary Note 10 for additional discussion). As a control, valinomycin-treated *B. subtilis* cells were stained with VF2.0.Cl and subjected to the same external K^+^ concentration changes. No significant variation in average lifetime <τ> was observed in these experiments (Supplementary Figure 4).

### Single-cell membrane potential estimation of *B. subtilis* using *V*_*m*_ *vs* <τ> calibration

We used the calibration curve obtained above to assess the membrane potential of *B. subtilis* cells in minimal medium (MSgg) in their unperturbed state based on the observed lifetimes (no valinomycin or added ions, Figure 6a, blue squares and Supplementary Table 4). Focusing first on the *mean* observed values, we found <τ>_MSgg_ = 845 ps on average (range: 772 – 920 ps), corresponding to a membrane potential *V*_*m*_,_MSgg_ = −64 mV (range: −99 to −49 mV). The largest value is similar to that observed for *B. subtilis* cells in MSgg treated with 25 µM valinomycin (Figure 6b, green squares): *V*_*m*_,_MSgg+Val_ = −50 mV (range: −53 to −43 mV). This value is smaller (in absolute value) than measured in M9 minimal media (see Supplementary Figure 6); <t>_M9_ = 734 ps on average (range: 620 – 804 ps), corresponding to a membrane potential of V_m,M9_ = −127 mV (range: −152 to −78 mV). This latter value is comparable to estimates reported by other research groups in different media and for different strains, which range from −80 to −140 mV (see Supplementary Note 11) (26-28,78). The differences between the MP measured in MSgg and M9, as well as those reported in the literature, highlight the effect of extracellular media composition, particularly the type and concentration of carbon and nitrogen sources, on the physiological state of bacterial cells, reflected in their membrane potential. MSgg, being a medium that promotes the growth of biofilms, imparts some physiological stress that results in a lower magnitude of membrane potential value. When membrane potential measured in a traditional minimal media M9, the magnitudes of membrane potential for B. subtilis matches quite well with the prior literature reports (26-28,78). The lifetime variation from series to series remained approximately constant across different conditions (Figure 6a-d), with mean lifetime dispersion in the “no perturbation” conditions of Figure 6a-b being comparable to that observed in high [K^+^]_out_ measurements (Figure 6c-d, orange and red squares). For these depolarized bacteria, we found <τ>_depol_ = 1,105 ps on average across different measurements (range: 1,075 – 1,210 ps) which translates into an average membrane potential V_m_^depol^= −14 mV (range: −19 – −1 mV).

**FIGURE 6:**
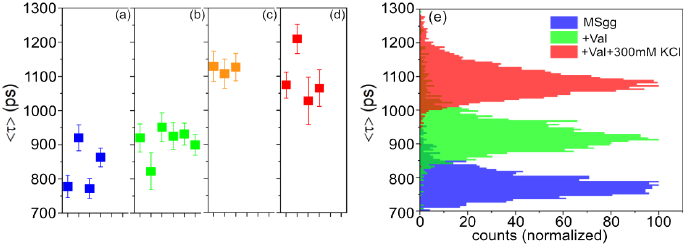
Average lifetime dispersion across measurements and external chemical conditions. Repeated measurements on different bacterial cultures resulted in slightly different measured average lifetimes in identical conditions. a: MSgg, b: MSgg + 25 μM valinomycin, c: same as (b) but with 240 mM KCl, d: same as (b) but with 300 mM KCl. e: In each measurement, the single-cell average lifetime was approximately normally distributed (with a few outliers not shown here), with standard deviation not fully accounted for by shot noise. Each histogram corresponds to the leftmost datapoint with the same color in the left panel (the histogram for (c) is omitted as it overlaps with that of (d)). MSgg: <τ> = 778 ± 32 ps, MSgg + Val: <τ> = 920 ± 41 ps, MSgg + Val + 300 mM KCl: <τ> = 1,075 ± 38 ps.

### Probing the effect of medium osmolality on membrane potential

The calibration measurements (Figure 4) and subsequent measurements performed at a different (constant) osmolality (Figure 5) used small ions (Na^+^, Cl^-^) to maintain a constant osmolality. While potassium is the dominant species in the equilibration of the membrane potential in the presence of valinomycin, it cannot be excluded that using the Nernst potential equation for K^+^ (Eq. (4)) instead of the Donnan equation (Eq. (3)) is leaving out some important details, especially at large external ion concentrations. In order to minimize these effects, we used xylose, a non-metabolizable and membrane-impermeable sugar to adjust the external medium’s osmolality (Figure 7, Table 1 and Supplementary Table 6).

**Table 1.**
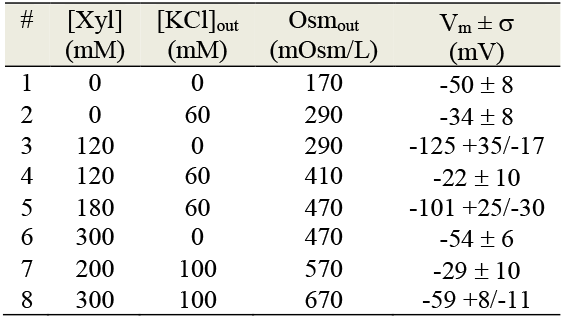
B. subtilis membrane potential measurements in different external concentrations of KCl and xylose.

**FIGURE 7:**
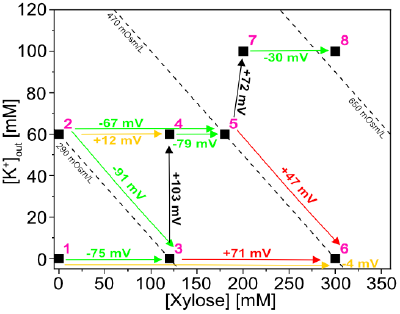
B. subtilis membrane potential measurements in different external concentrations of xylose and KCl in the presence of 25 μM of valinomycin. The black squares indicate the external xylose and KCl concentrations to which the cells were exposed. The measured Vm for the different measurements are tabulated in Table 1. Some of these conditions differ only by their [K+]out, while some others differ only by their osmolality (∼[Xylose] + 2[K+]out). The green and red arrows connect conditions between which either Δ[K+]out = 0 and ΔOsm ≠ 0, or ΔOsm = 0 and Δ[K+]out ≠ 0. The colors indicate agreement (green) or disagreement (red) with the sign of the membrane potential variation predicted by the Nernst potential (see Supplementary Table 6). The change in Vm is indicated next to each arrow. The orange arrows correspond to cases where almost no change in Vm is observed (within the typical measurement uncertainty of ± 20 mV), consistent with either a positive or negative change. The black arrows indicate changes where Δ[K+]out ≠ 0 and ΔOsm ≠ 0 but the sign of change cannot be predicted from the Nernst potential. Equi-osmolality lines are indicated by dashed diagonal lines.

Increasing the medium osmolality at constant [K^+^]_out_ should result in water expulsion from the cell in order to balance the osmotic pressure. Consequently, an increase of [K^+^]_in_ is expected, as has been well documented in *E. coli* (80). This behavior, and the internalization of permeable solutes in the cytoplasm, is a general osmo-adaptation strategy of bacteria in response to physiological changes (81). Assuming that water efflux dominates, this should be followed, according to the Nernst potential (Eq. (4)), by a decrease of the membrane potential. Similarly, increasing [K^+^]_out_ while keeping the medium osmolality constant should have little effect on [K^+^]_in_, and therefore, according to Eq. (4), should be followed by a membrane potential becoming less negative. Figure 7 indicate these two types of changes by horizontal (Δ [K^+^]_out_ = 0, ΔOsm_out_ > 0) or diagonal (Δ [K^+^]_out_ > 0, ΔOsm_out_ = 0) arrows, their colors indicating agreement (green) or disagreement (red) with the above prediction of the Nernst potential. A third type of change is indicated by black vertical arrows in Figure 7, corresponding to cases where Δ[K^+^]_out_ > 0 and ΔOsm > 0, the increase in osmolality being mostly due to increasing [K^+^]_out_.

Under these conditions, two opposing effects happen simultaneously, making the Nernst equation useless to predict the sign of the expected membrane potential change. Experimentally, our observations suggest that in conditions of large osmolality, the dominating effect is that of Δ[K^+^]_out_ > 0, associated with an increase of the membrane potential. The specificity of these stressful conditions could explain why our naïve use of the Nernst equation fails to predict the correct sign for the membrane potential change in this regime (red arrows in Figure 7), in particular if K^+^ influx is involved (82).

### Color-coded visualization of the membrane potential in *B. subtilis* cells

In the previous discussions, we have estimated average lifetime values (and hence membrane potential) from individual cells (utilizing integrated signals). However, thanks to the independence of phasor calculations from SNR, it is formally possible to compute an amplitude-averaged lifetime for each individual pixel in the image, which can be used to obtain color-coded lifetime maps (Figure 2c) or membrane potential maps. Given their lower SNR, the phasors of individual pixels are expected to exhibit a larger dispersion than those of individual cells, Consequently, the single-pixel amplitude-averaged lifetimes are expected to be affected by a larger uncertainty than those of individual cells, and so is that of the membrane potential thus obtained. The calibration curve of Figure 4c has been used to convert single-pixel amplitude-averaged lifetimes into membrane potential, which then color-coded and overlaid on the original image (Figure 8 and Supplementary Figure 5). Figure 8 clearly shows that the unperturbed *B. subtilis* cells in minimal medium (MSgg) are characterized by a larger negative membrane potential (Figure 8a and Supplementary Figure 5) than the cells exposed to a depolarizing chemical condition such as a high concentration of KCl in the presence of valinomycin (Figure 8b). These cells exhibit a close to zero membrane potential, as expected from the loss of K^+^ gradient across the membrane. Membrane potential maps enable visualizing not only the cell-to-cell variability observed in all single-cell results discussed in this work, but potentially as well, spatial inhomogeneity within individual cells irrespective of extracellular chemical conditions. However, sub-cellular spatial inhomogeneity of MP is difficult to characterize as the single-pixel SNR is insufficient. A majority of pixels had counts of *N* = 100 photons or lower, leading to an average lifetime uncertainty of approximately 146 ps or 30 mV. (see Supplementary Note 5 for details.) Future work otpimizing SNR and imaging conditions will look into this intriguing potential of our bacterial MP measurement approach.

**FIGURE 8.**
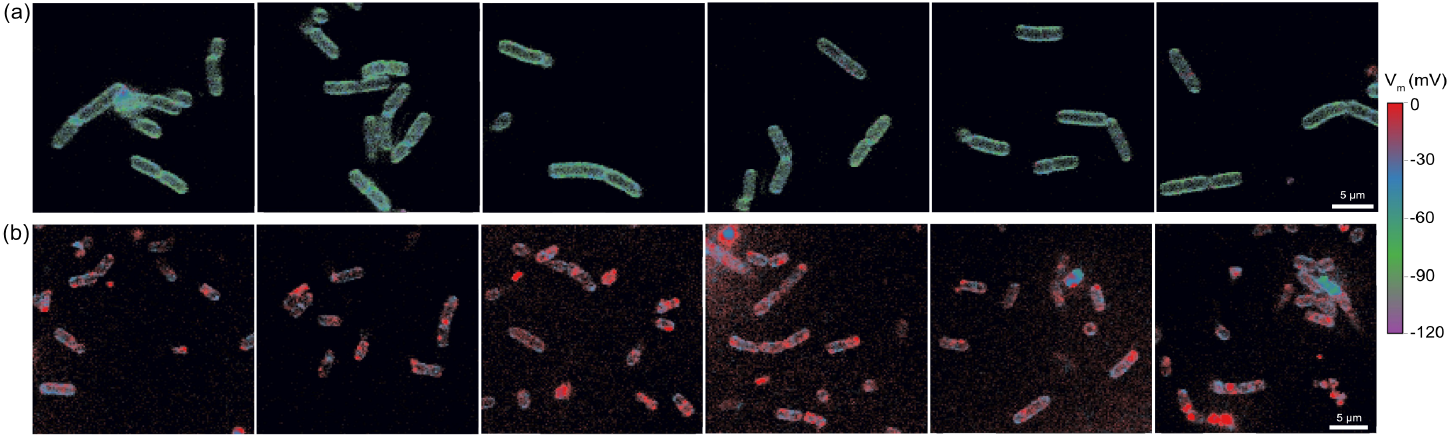
Effect of varying extracellular chemical conditions on the membrane potential of B. subtilis. (a) Top panel: Unperturbed cells in MSgg only: Vm ∼ −60 mV, (b) Bottom panel: Chemically depolarized cells in MSgg (25 µM valinomycin + 240 mM KCl): Vm ∼ −14 mV. Each box in this figure represents a cropped-in ROI of ∼ 27.5 µm x 27.5 µm, taken from images acquired from a field of view (FOV) of ∼ 110 µm x 110 µm. See Supplementary Figure 5 for the full FOV average lifetime and membrane potential images. Scale bar = 5 µm.

## Conclusion & outlook

In conclusion, we have explored the use of a novel kind of voltage-sensitive dye, VF2.1.Cl, for the study of membrane potential changes in a model Gram-positive bacterium, *Bacillus subtilis* for the first time. To calibrate the fluorescence response of VF2.1.Cl to changes in the membrane potential of *B. subtilis*, we altered the potassium concentration gradient across the cell membrane using a known ionophore, valinomycin. We obtained a nonlinear relation between amplitude-averaged VF2.1.Cl fluorescence lifetime and membrane potential. We have thus demonstrated the potential of VF2.1.Cl as a fluorescence lifetime indicator for *B. subtilis*’s membrane potential changes. We have taken advantage of the simplicity of phasor analysis to analyze VF2.1.Cl’s fluorescence lifetime at the single-cell level, and to rapidly represent membrane potential information at the single pixel level in FLIM images containing several cells. In particular, we have introduced a simple internal population-level approach to define single-exponential references used in the phasor analysis of the observed biexponential fluorescence decays. We estimated the membrane potential of unperturbed *B. subtilis* cells to be 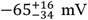 (in MSgg) and 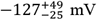 (in M9), while chemically depolarized cells exhibited a membrane potential of 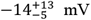. Beyond simply estimating membrane potential, we have characterized and quantified the shot noise contribution to the observed lifetime dispersion. The observation of a ‘non-shot noise’ residual ∼30-50 ps lifetime standard deviation, indicates the existence of ∼6-10 mV of MP variability within individual cells of a *B. subtilis* populations. This MP variability across individual cells likely arises from differences in metabolic and physiological states between individual cells.

In summary we have developed methods that allows bacterial MP changes to be easily visualized, recorded and quantified. Our measurements allowed interpreting most of the observed variations in VF2.1.Cl lifetime (and hence membrane potential) upon media (chemical composition) changes in a way that appeared for the most part consistent with the K^+^ Nernst potential approximation for the membrane potential, and a simple interpretation of the effect of osmotic pressure change on the bacterial volume. The possibility to determine the membrane potential at localized regions of a cell membrane is a significant advantage of this method over other techniques. The potential applications of this capability are vast and will be explored in future work.

Our work raises a number of questions which further experiments will need to address. The first concerns the differences in measured lifetimes of VF2.1.Cl in the membrane of *B. subtilis vs* those observed in eukaryotic cells. The range of lifetimes measured in *B. subtilis* cells (suspended in MSgg) with MP ranging from ∼ −100 mV to ∼ −10 mV was ∼ 800 ps to 1.2 ns, significantly shorter than the lifetimes previously measured in eukaryotic cells (*e*.*g*. ∼ `1.5 ns ≤ <τ> ≤ 2.1 ns in HEK293T cells with −80 mV < *V*_*m*_ < +80 mV (38)). This is related to the different decay constants measured in these two systems. While Lazzarri *et al*. reported lifetime components *τ*_*1*_ = 2.6 ns and *τ*_*2*_ = 0.9 ns, we observed *τ*_*1*_ in the [2.4, 3.2] ns range (median: 2.76 ns) and *τ*_*2*_ in the [350, 800] ps range (median: 550 ps) depending on medium composition. This contributes to the shorter average lifetimes observed in this work. Lazarri *et al*. reported a decrease in average lifetimes at high dye concentrations, but the dye concentration used in our work are in the low end of the range they explored, making dye concentration an unlikely explanation of these lifetime differences in bacteria and mammalian cells. However, bacterial cytoplasmic membranes are quite different from eukaryotic ones in their structure and composition (83,84). Bacterial membranes are primarily composed of phospholipids and lack rigidifying sterols for instance, which makes a bacterial cell membrane more fluidic than a mammalian cell membrane. In particular, bacterial membrane can respond to environmental changes such as temperature, pH and osmotic pressure rapidly by altering their lipid composition (85-87). Additionally, the *B. subtilis* membrane possesses a ∼34 nm thick external porous peptoglycan wall separating it from the external medium (88), which, may provide a different chemical environment for the membrane bound probe. As discussed in Supplementary Note 8, the charge distribution within the bacterial cell wall could generate a significant electrical potential gradient outside the bilayer. This gradient could influence the effective potential drop across the membrane, which is detected by the dye (89,90). In essence, all these factors can potentially contribute to the observed differences in VF2.1.Cl’s distinct photophysical characteristics.

The second and related question concern the voltage sensitivity of VF2.1.Cl in *B. subtilis*. In this work, we found that the average lifetime of VF2.1.Cl changes by ∼5 ps/mV in the range −60 mV ≤ V_m_ ≤ 0 mV, its sensitivity dropping to ∼1.2 ps/mV for more negative V_m_ values. This is different from the 3.5 ps/mV over the [-80, 80] mV range reported in HEK293T cells reported by Lazarri *et al*. Because this sensitivity is related, among other factors, to the respective orientation of the electric field and electric transition dipole of the molecule (supplementary Eq. S8) as well as their magnitude, this could indicate a different electric field magnitude in the bacterial membrane (due to a different membrane thickness, or as mentioned above, a contribution of the cell wall to the electric potential gradient), or a different average molecular orientation of VF2.1.Cl in the membrane.

Third, there exists large uncertainty in estimated membrane potential values for lower membrane potential value beyond ∼ −80 mV. Lower membrane potential (below ∼ −80 mV)/smaller lifetime (below ∼ 800 ps) values correspond to physiological conditions where according to Nernst equilibrium a small change of external potassium concentration leads to significant changes in the membrane potential (as illustrated in Figure 4a). Thus, any uncertainty in the estimation of lifetime leads to magnified uncertainty in the estimated potential during the conversion process. Interestingly, the uncertainty on lifetime estimations at low external K+ concentration (as evident in Figure 4b) is comparable to that observed in conditions of larger external K+ concentration, at equivalent fluorescence intensity. But there exists a significantly larger contribution of population variability (as can be seen from the reduced contribution of shot noise to the observed dispersion, see Supplementary Table 5). This is likely due to the greater biochemical stress experienced by the cells. The dependence on membrane potential on [K^+^]out is an inherent biological phenomenon that cannot be eliminated. Nevertheless, the second source of uncertainty could potentially be reduced by performing higher SNR measurements. This would require longer cell exposures and could potentially result in increased cell stress, further amplifying the population dispersion of the observed membrane potential.

Fourth, the fact that our calibration approach relies on past work that is difficult to reproduce, and that we observed differences between measurements in presumably identical conditions (e.g. Figure 5), hint at sources of variability that were not fully under control. Our work should thus be considered a first step towards establishing a robust *V*_*m*_ *vs* <τ> calibration for VF2.1.Cl in *B. subtilis*. One way to better control the cells’ physiological state and minimize the delay between harvesting, medium replacement, staining and observation could involve studies performed in temperature-controlled microfluidic chambers, allowing rapid *in situ* medium exchange. A similar technology would also enable monitoring individual cell membrane potential evolution over time, whether isolated or within biofilms.

Alternatively, more direct ways of modulating the cell membrane potential such as induced transmembrane voltage application could be used, although it is unclear how reliably the actual applied field can be estimated based only on a geometric model of the bacterial shape (24,25).

Notwithstanding, these open questions, which we plan to address in future work, our current results demonstrate the exciting potential of phasor-based, fluorescence lifetime measurement of VF dyes for precise and absolute potential measurements in bacterial cells. We anticipate that these and similar molecular indicators, together with advances in wide-field time-resolved detectors enabling simpler FLIM measurements (61,91), will enable new insights in the complex bioelectrical activity of bacterial communities.

## Materials & Methods

### Sample preparation for imaging

*B. subtilis* culture, growth and membrane staining protocols are described in Supplementary Note 1. Aliquots of *B. subtilis* cells were taken at the mid-log phase (during subculture) and were incubated with VF2.1.Cl for ∼30 mins for staining. After successful staining and washing, the stained cells were divided into two groups. One group (a) was left unperturbed, while the other group (b) was subjected to different extracellular chemical environments. Group (b) was treated with Valinomycin, KCl, NaCl, or xylose to mimic artificial modulation of MP. Stained bacterial cells were suspended in poly-L-lysine coated Petri dish/IBIDI flow channels with modified minimal salts glycerol glutamate (MSgg) medium, an optically transparent minimal medium (see Supplementary Table 1). Valinomycin, KCl, NaCl or xylose were added directly to the MSgg medium and the cells left incubating for at least 30 minutes before imaging. The VoltageFluor does not exhibit any growth-inhibitory effects. the cells continue to grow and divide after staining with voltage probe VF2.1.Cl (see Supplementary Note 3). We have also tested for growth inhibitory effects of imaging/exposure to laser irradiation. The cells continue to grow after repeated laser irradiation (see Supplementary Note 3).

### Fluorescence lifetime imaging microscopy (FLIM)

Several fields-of-view (FOV) of stained bacterial cells were imaged with a laser scanning confocal microscope equipped with time-correlated single-photon counting (TCSPC) hardware for fluorescence lifetime imaging microscopy (FLIM) (detailed in Supplementary Note 2).

Fluorescence lifetime information was analyzed using AlliGator, our freely available phasor- and NLSF FLIM analysis software, allowing cell-level or pixel-level fluorescence decay analysis (59,63),(92).

During our investigations into membrane potential variations in response to different extracellular chemical conditions and calibration measurements, we have analyzed the data by treating each individual cell as a single entity. We have used integrated signals from individual cells to estimate lifetime of each cell, thereby determining it’s membrane potential, regardless of its morphological characteristics. All pixels within the contour of each individual cell (including the fluorescence inside the cell, which, as argued in the text and Supplementary Note 7, corresponds to out-of-focus fluorescence from the top and bottom of the cell) are considered for lifetime analysis.

### Fluorescence lifetime estimation by NLSF

In some cases, fluorescence lifetime estimation was done by nonlinear least-square fitting of the recorded photon arrival time histogram. The measured fluorescence decay can be expressed as (63):

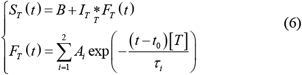

where the notation introduced in ref. (62) are used, in which an index *T* indicate a *T*-periodic function (*T* being the laser period). The observed signal *S*_*T*_(*t*) is the convolution of the instrument response function (IRF) *I*_*T*_ with the fluorescence decay *F*_*T*_(*t*), modeled as a *T*-periodic bi-exponential function, plus a baseline parameter *B* accounting for residual uncorrelated background and an offset *t*_*0*_. *t*[*T*] denotes *t* modulo *T. A*_*i*_ and *τ*_*i*_ are the amplitude and lifetime of component *i*. where the amplitude *A*_*i*_ is related to signal intensity *N*_*i*_ (number of photocounts) by *N*_*i*_*= A*_*i*_*τ*_*i*_. The amplitude fractions are defined as *α*_*i*_ =*A*_*i*_/(*A*_1_+*A*_2_) from which the amplitudeaveraged lifetime ⟨ *τ* ⟩_*a*_ is given by:

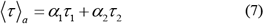

while the intensity fractions are defined as *f*_*i*_ =*N*_*i*_/(*N*_1_+*N*_2_). Parameters *B, A*_*1*_, *A*_*2*_, *τ*_1_. and *τ*_2_ are obtained using a standard iterative Levenberg-Marquard algorithm to minimize the (Poisson) weighted *χ*^*2*^ function (63).

### Phasor analysis of fluorescence lifetime

Decays were converted into a calibrated phasor (*g, s*) as the normalized components of the first term of the decays’ Fourier series as previously described (59), using the IRF as calibration associated with lifetime *τ* = 0 ns. Single pixel or single region of interest (ROI) phasors can be represented as a 2D histogram in the (*g, s*) plane. The phasor plot provides a user-friendly representation of complex decays due to its unique properties. Phasors corresponding to single exponential fluorescence decays are located on the so-called universal (semi)circle (UC) of radius 1/2 centered on (*g, s*) = (1/2, 1/2). In the case of a biexponential decay, the phasor is located on the segment connecting the two single-exponential decays’ phasors (or “references”). References *τ*_*1*_ and *τ*_2_ were defined internally for each condition, by plotting a scatterplot of all cell-level phasors in that condition and finding the intersection of the major axis of inertia of the scatterplot with the universal circle (UC), location of single-exponential decays in the phasor representation (Figure 3b). This approach of determining individual single exponential reference lifetime components is a first of it’s kind.

The relative distances of the phasor to these references (or phasor ratios, geometric quantities that are easily computed once the phasors are computed) correspond to the relative fractions *f*_*i*_of each normalized single-exponential’s phasor and are identical to the intensity-fractions of the single-exponential components of the decay (59).

From these fractions, a mean lifetime can be computed as (59):

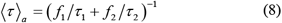

This phasor ratio-based average lifetime is formally identical to the amplitude-averaged lifetime defined by Eq. (7) (59). All mention of average lifetimes in the remainder of the text will refer to either one of these equivalent definitions of the amplitude-averaged lifetime, as it is the one relevant for the analysis of membrane potential with VoltageFluor dyes, as discussed in the next section. The index *a* (for amplitude-averaged) will also be dropped for simplicity.

A brief description of some of the phasor analysis steps follows, more details being provided in Supplementary Note 4.

### Color-coded average lifetime map

In order to visualize individual bacterial cell MPs under different extracellular chemical conditions we have created color-coded lifetime maps and hence membrane potential maps. Amplitude-averaged lifetime were computed for each individual pixel within a cell using Eq. (8). A color scale was then used to encode amplitude-averaged lifetimes within a user-selected range, color which was then overlaid on the original image, the original pixel intensity (contrast-adjusted for clarity) being used to set the color brightness, as illustrated above in section 8.

### Estimation of shot noise influence on the amplitude-averaged lifetime

In order to assess the relative contributions of population heterogeneity and signal-to-noise ratio (SNR) to the average lifetime distributions observed in our experiments, we performed Monte Carlo estimations of the influence of shot noise (the main noise contribution in the TCSPC measurements performed here) to the computed single-cell average lifetime in the following manner. For each cell fluorescence decay {*I*(*t*_*i*_)}_*i=1*,*…*,*G*_, we simulated *N* = 1,000 replicas of this decay, replacing each *I*(*t*_*i*_) by a random number Poi(*I*(*t*_*i*_)), and computing the corresponding <τ> based on the same phasor references. The corresponding standard deviation σ_<τ>_^SN^ of the distribution of average lifetime is an estimation of the uncertainty on <τ> computed for the single-cell decay due to shot noise, an approximation supported by direct simulation of shot noise-limited decays with identical photon counts (detailed in Supplementary Note 5).

## ASSOCIATED CONTENT

## Supporting Information

A Supplementary Information file containing additional figures, tables and notes is available free of charge on the Biophysical Reports website. Supplementary figures depict comparison of fluorescence decays and average lifetime for VF2.1.Cl and VF2.0.Cl in various conditions, effect of increasing [K^+^]_out_ on average lifetime for VF2.1.Cl in a cell, average lifetime and membrane potential maps for different fields of view corresponding to unperturbed and chemically depolarized chemical conditions, average lifetime dispersion across measurements. Supplementary tables include data for MSgg composition, VF2.1.Cl and VF2.0.Cl fluorescence decay fit parameters in various conditions, average lifetimes for VF2.1.Cl in a cell under various extracellular chemical conditions, shot noise contribution to the total standard deviation of the average lifetime. Supplementary notes provide details on cell culture, growth and membrane staining protocols, FLIM data and brightfield image acquisition, effect of imaging conditions on B. subtilis cell growth, additional details for phasor analysis, shot noise influence on phasor-based amplitude-averaged lifetime, 3-state model of the photophysics of VoltageFluor dyes, fluorescence intensity profile analysis for stained bacterial membranes, estimation of the potential drop across a semipermeable membrane in the presence of a membrane potential, additional calibration analysis details, fluorescence lifetime measurements outside cells, literature reporting estimated membrane potential values for B. subtilis and ‘puncta’ in membrane labelling.

## Data availability

Raw data files and analysis results will be freely available on FigShare by the time of publication.

## Software availability

AlliGator is freely available on Github.(92)

## AUTHOR INFORMATION

## Author Contributions

DR, XM & SW designed the experiments. DR, KB performed experiments and DR conducted data analysis. XM wrote the software, theory and contributed data analysis concepts. EM provided the VFs. DR and XM wrote the manuscript, which was edited by all other authors. All authors have given approval to the final version of the manuscript.

## Notes

The authors declare no competing financial interest.

