## Supplemental Information for "Towards measurements of absolute membrane potential in Bacillus subtilis using fluorescence lifetime"

This document contains supplementary figures, tables and notes referred to in the main text.

### Supplementary Figures

#### Supplementary Figure 1: Observed VF2.0.Cl and VF2.1.Cl fluorescence decays in different conditions

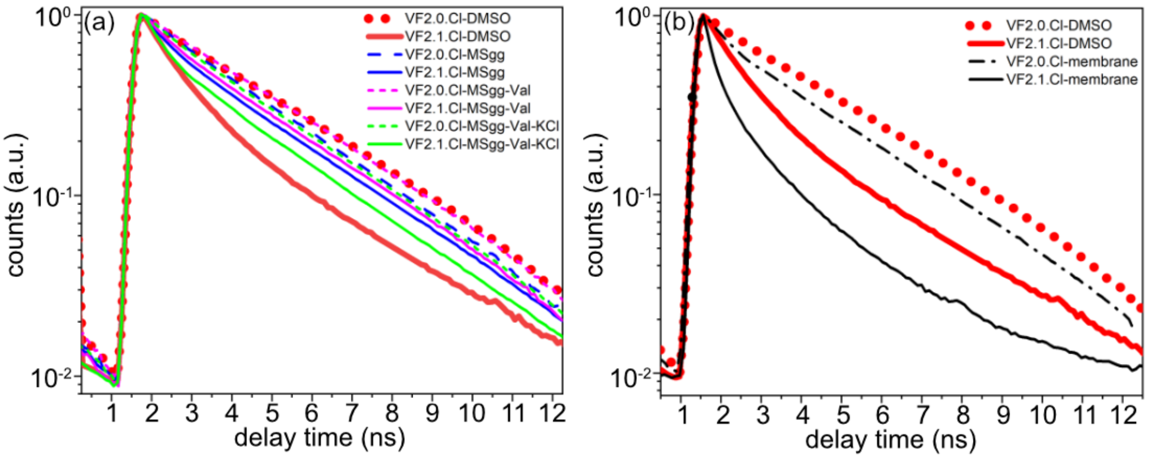

**Supplementary Figure 1**: (a) Comparison of fluorescence decays for VF2.1.Cl and VF2.0.Cl in various solutions (without cell). (b) Comparison of fluorescence decays for VF2.1.Cl and VF2.0.Cl in DMSO and in live *B. subtilis* membrane.

#### Supplementary Figure 2: Effect of increasing $[K^+]_{out}$ on average lifetime of VF2.1Cl in cell

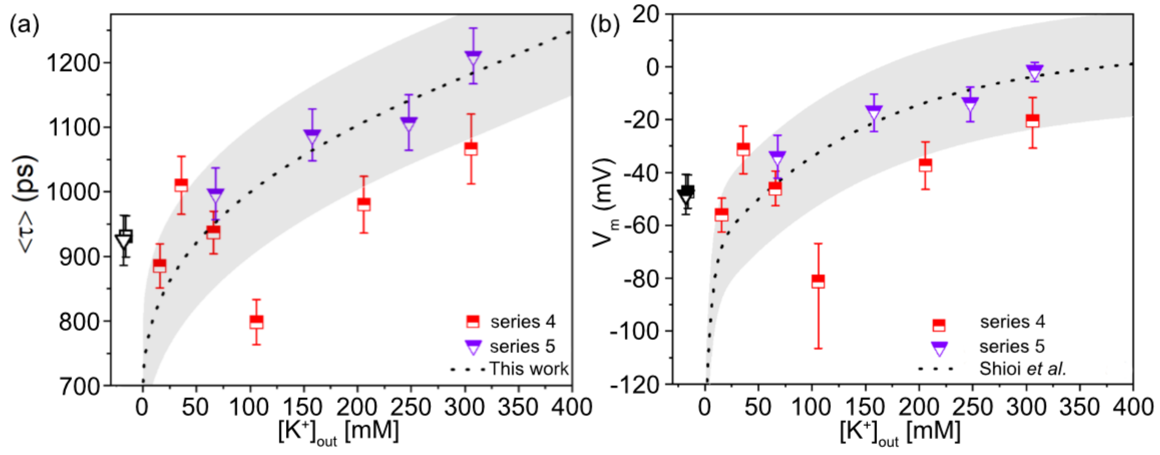

**Supplementary Figure 2**: Effect of increasing $[K^+]_{out}$ on <τ>. *B. subtilis* cells in the mid-exponential phase were treated with 25 µM valinomycin and supplemented with various concentrations of KCl (1 to 300 mM). These experiments were performed without maintaining constant osmolality. In practice, the medium osmolality increased from ~180 to 770 mOsm/Lwhen increasing $[K^+]_{out}$ from 5 to 300 mM. (a) $\langle\tau\rangle \pm \sigma_{\langle\tau\rangle}$ as a function of $[K^+]_{out}$ for two different repeats of the same measurement. The black data points correspond to measurements in MSgg only and are artificially offset on the $[K^+]_{out}$ axis for legibility purpose. The dashed line corresponds to the values measured in the calibration series (obtained at constant external osmolality) and is provided as a guide to the eye. The grayed area corresponds to the calibration curve values ± 100 ps. (b) Corresponding membrane potential values calculated using the calibration curve established in this work. The dashed line corresponds to the membrane potential values measured in the work of Shioi et al. (at constant external osmolality). The grayed area corresponds to these values ± 20 mV.

#### Supplementary Figure 3: VF2.1.Cl average lifetime in different conditions

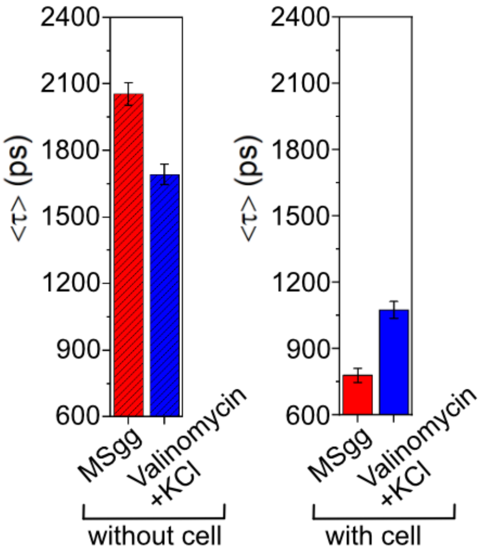

**Supplementary Figure 3:** Effect of valinomycin and KCl on VF2.1.Cl in MSgg medium and in the membrane of B. subtilis cells. In the absence of B. subtilis cells, VF2.1.Cl dissolved in MSgg with 25 µM valinomycin and 300 mM KCl exhibits a decreased $\langle\tau\rangle$ compared to pure medium (MSgg only). Note that both average lifetimes are larger than that of VF2.1.Cl in DMSO ($\langle\tau\rangle_{2.1,DMSO}$ = 1.53 ns). We do not have an explanation for these solvent effects. When VF2.1.Cl is loaded in the membrane of B. subtilis, the observed average lifetime decreases due to the presence of a large negative membrane potential. In the presence of 25 µM valinomycin and 300 mM KCl, the membrane is depolarized, resulting in an increased $\langle\tau\rangle$. The concentration of VF2.1.Cl used in all experiments was ~100 nM. In the case of B. subtilis measurements, the free dye was washed away before measurement.

#### Supplementary Figure 4: VF2.0.Cl average lifetime in different conditions

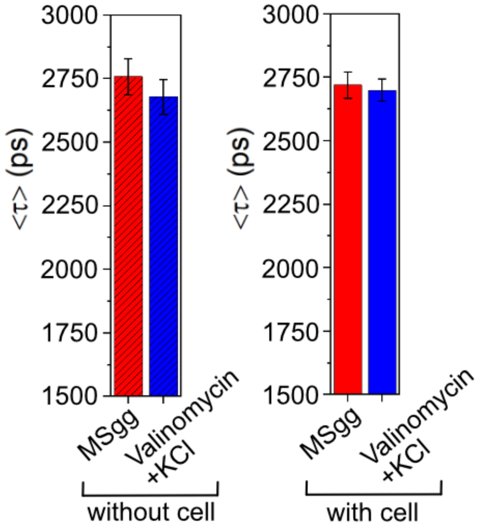

**Supplementary Figure 4**: Effect of valinomycin and KCl addition on VF2.0.Cl's lifetime in MSgg medium and in the membrane of *B. subtilis* cells. When VF2.0.Cl is in MSgg without cells, direct interaction with 25 µM valinomycin and 300 mM KCl does not result in significant changes of <τ> in contrast to VF2.1.Cl (see Supplementary Figure 3). Note that both average lifetimes are slightly *smaller* than that of VF2.0.Cl in DMSO ($\langle\tau\rangle_{2.0,DMSO}$ = 3.21 ns). We do not have an explanation for these solvent effects on VF2.0.Cl's lifetime at this stage of our work.

When VF2.0.Cl is in the membrane of *B. subtilis*, 25 µM valinomycin and 300 mM KCl do not cause significant changes in <τ> unlike what is observed with VF2.1.Cl (see Supplementary Figure 3). Similar concentrations of VF2.0.Cl (~100 nM) were used in all experiments. Interestingly, <τ> for VF2.0.Cl in solution and in the bacterial membrane are slightly different, suggesting a sensitivity of the fluorophore to the membrane environment not related to PeT.

#### Supplementary Figure 5: Average lifetime and membrane potential maps for different fields of view corresponding to unperturbed and chemically depolarized chemical conditions.

FOV1: unperturbed, in MSgg only

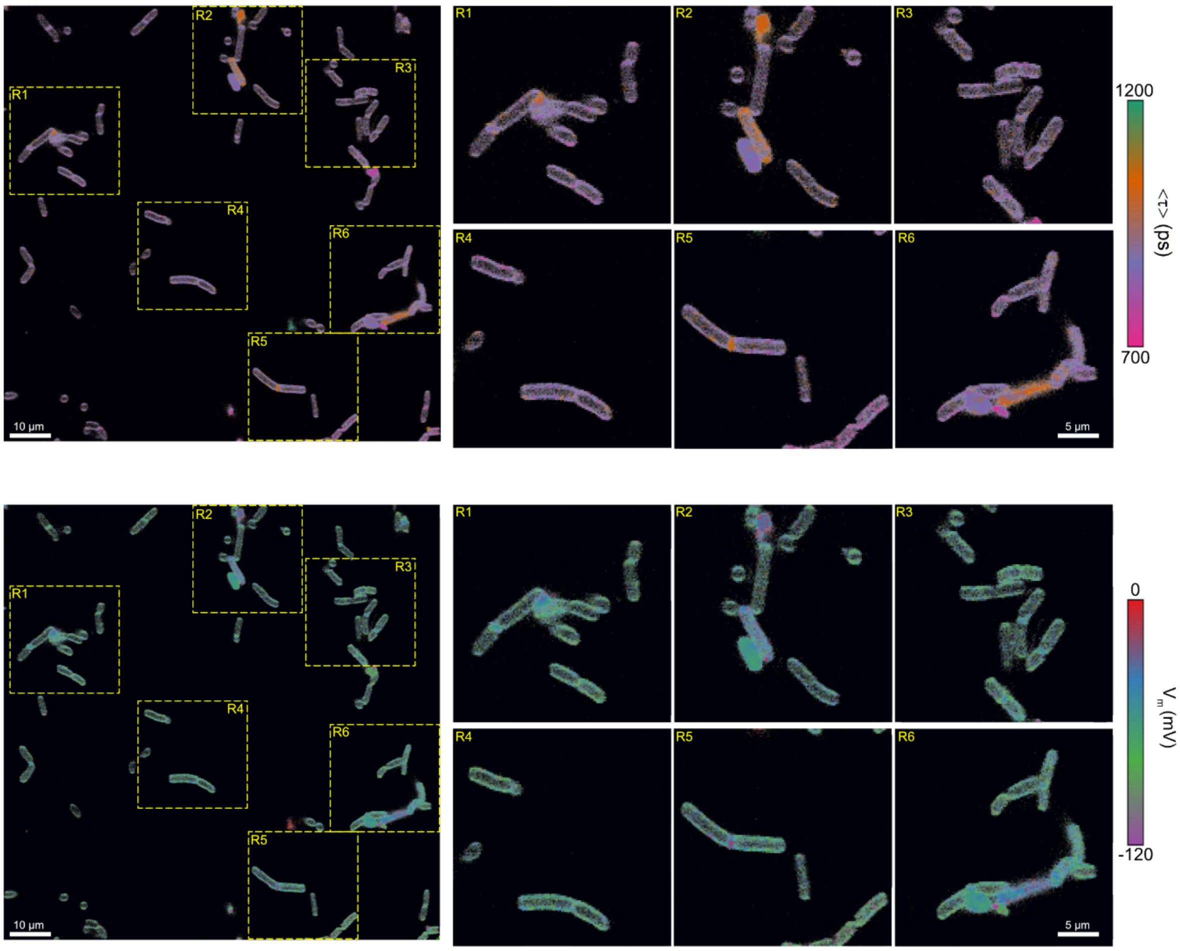

FOV2: unperturbed, in MSgg only

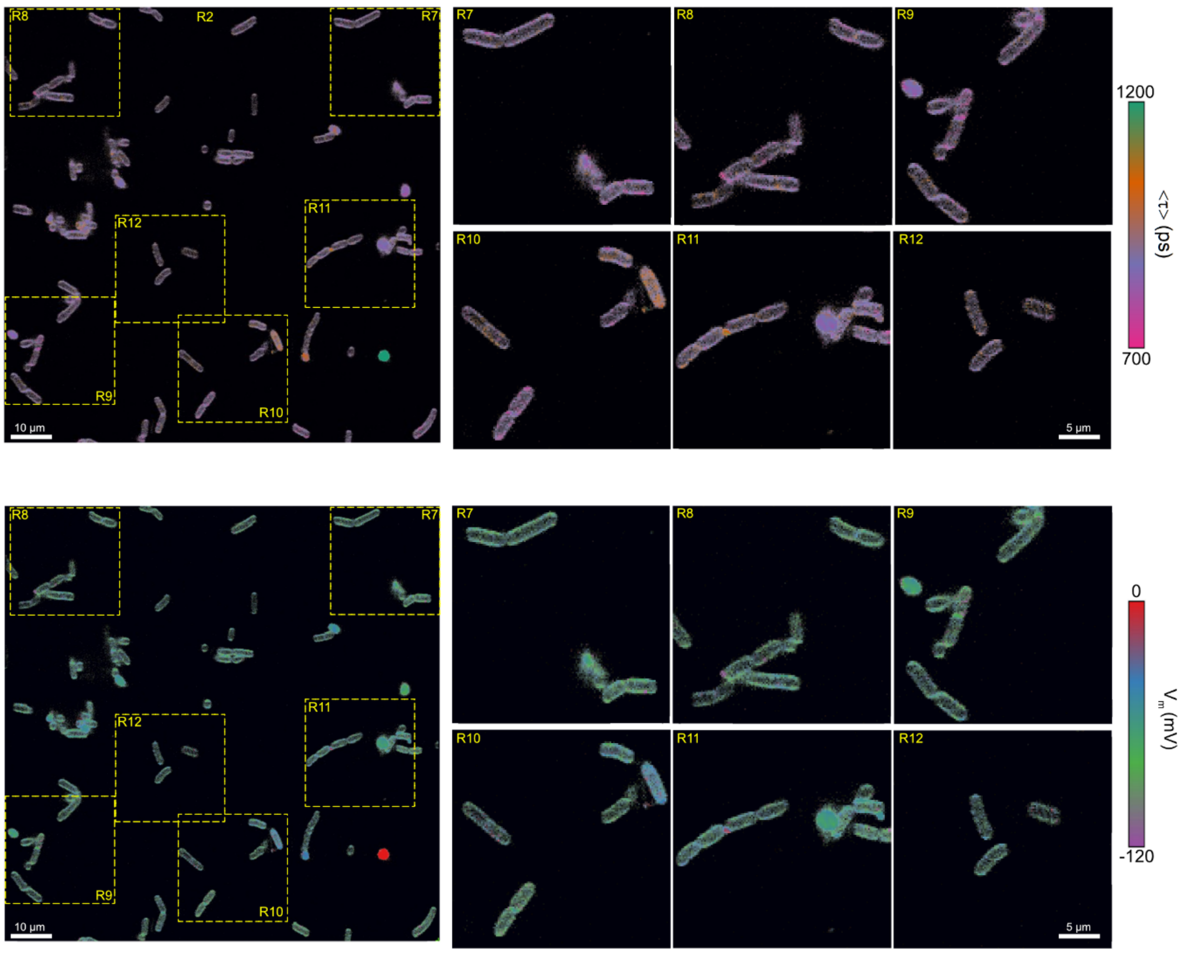

FOV3: unperturbed, in MSgg only

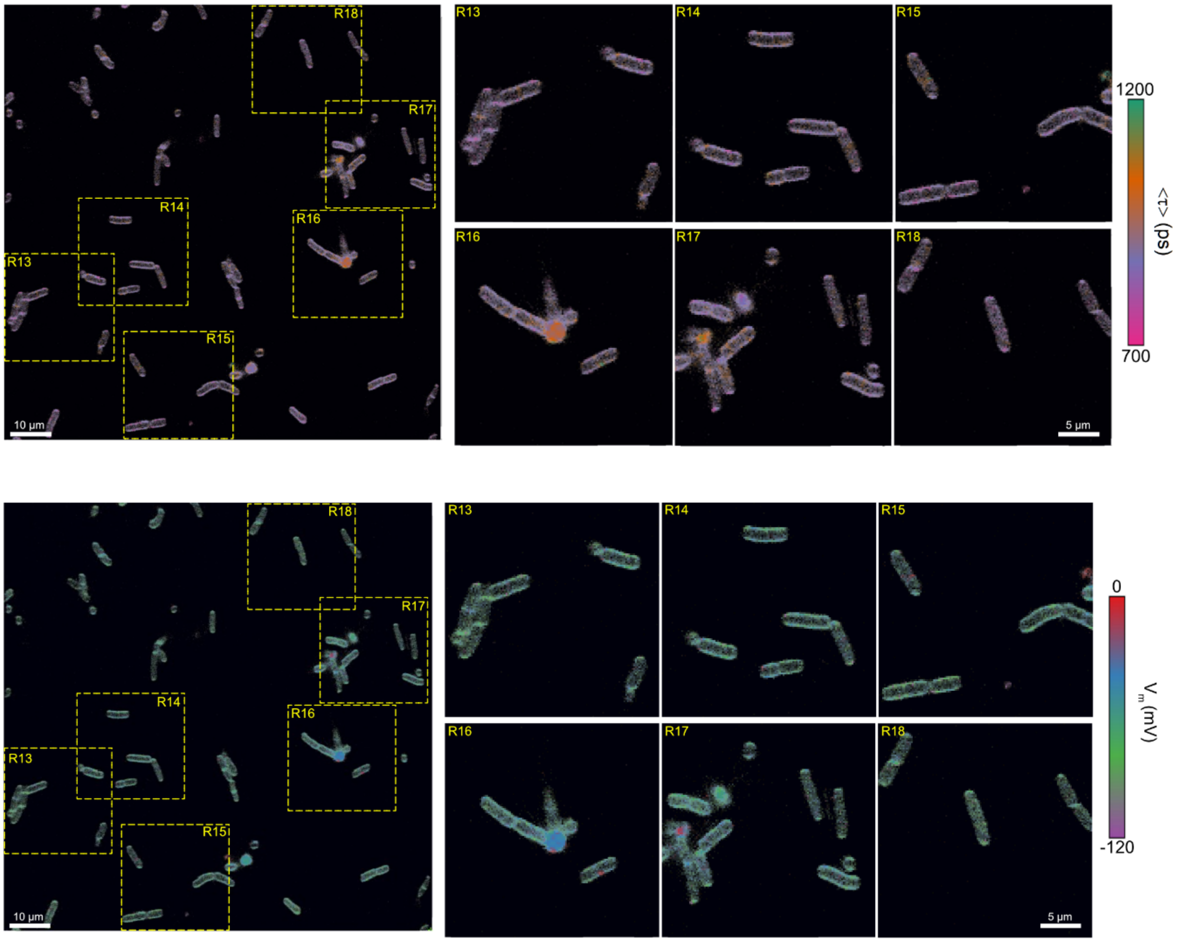

FOV4: chemically depolarized, treated with 25μM Valinomycin and 240 mM KCl

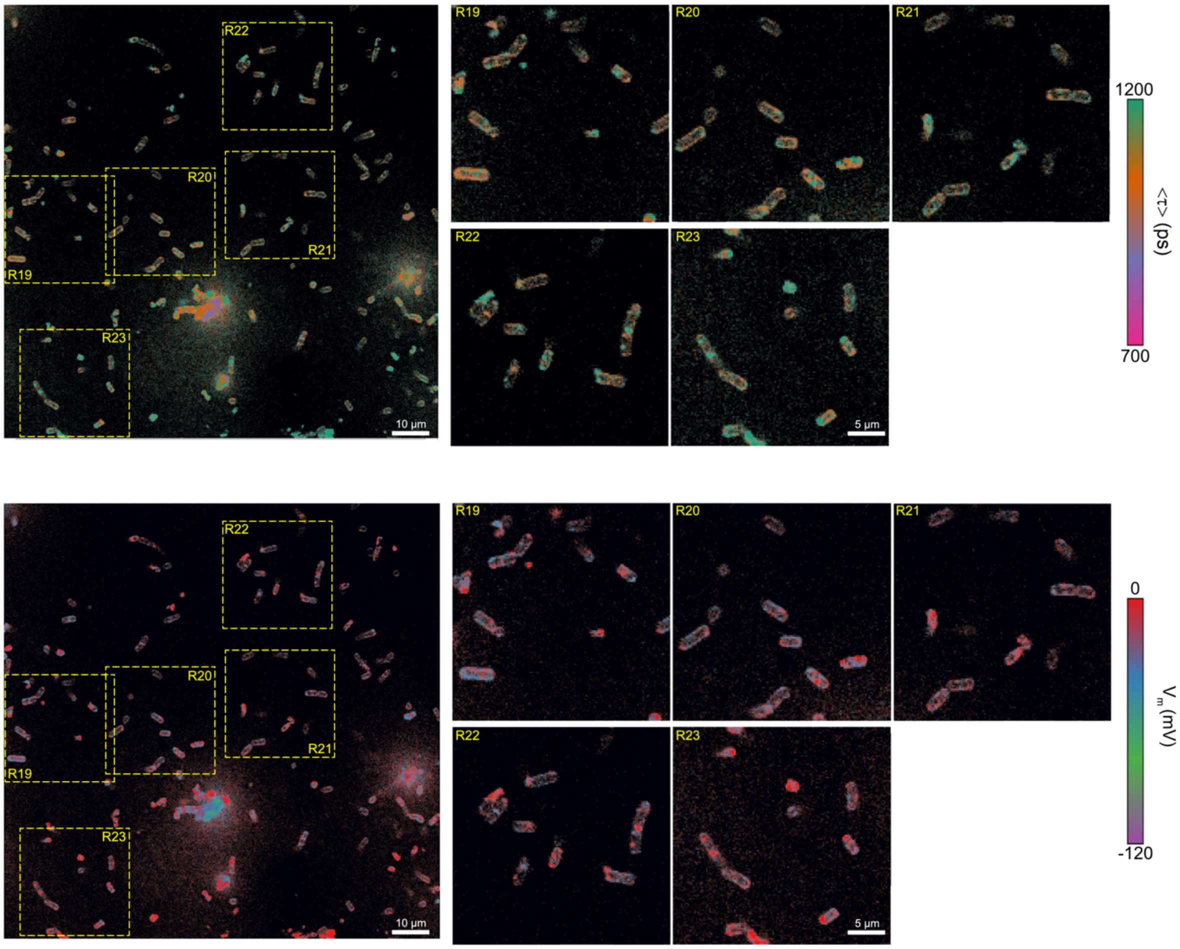

FOV5: chemically depolarized, treated with 25µM Valinomycin and 240 mM KCl

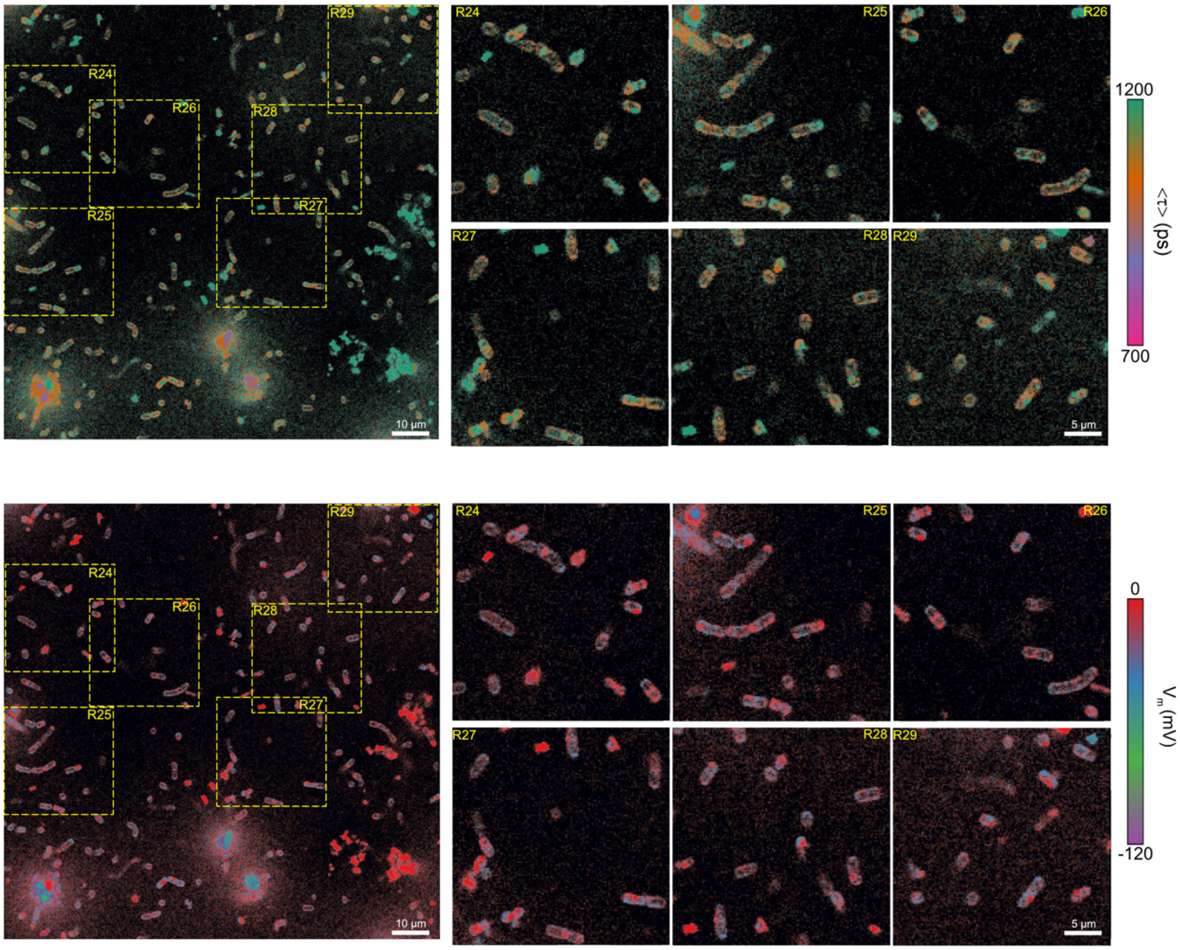

FOV6: chemically depolarized, treated with 25µM Valinomycin and 240 mM KCl

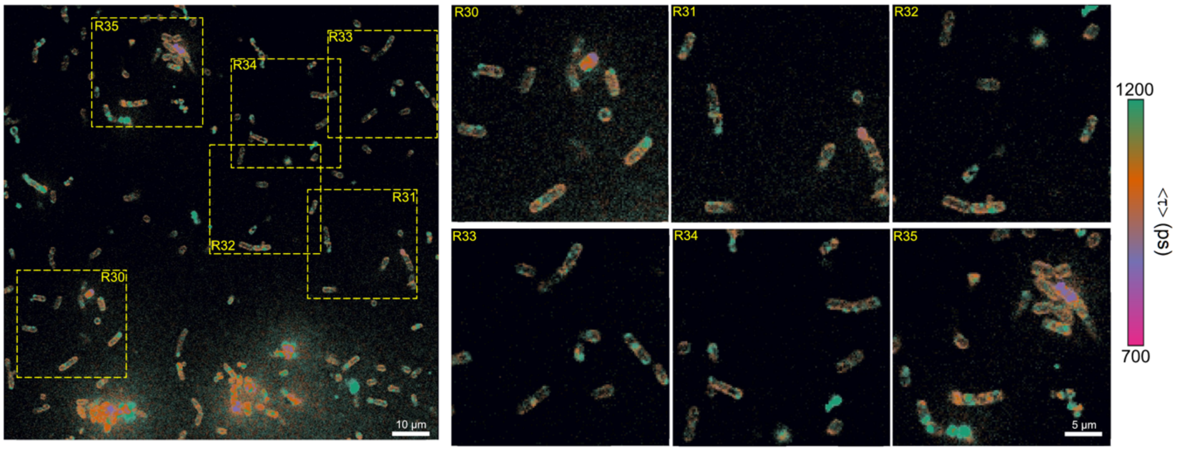

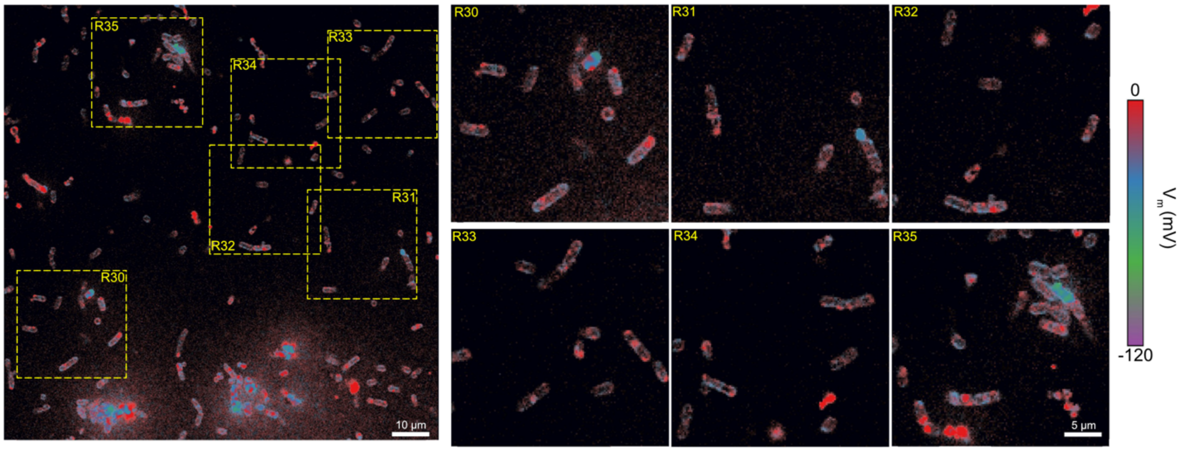

FOV7: unperturbed, in M9 only

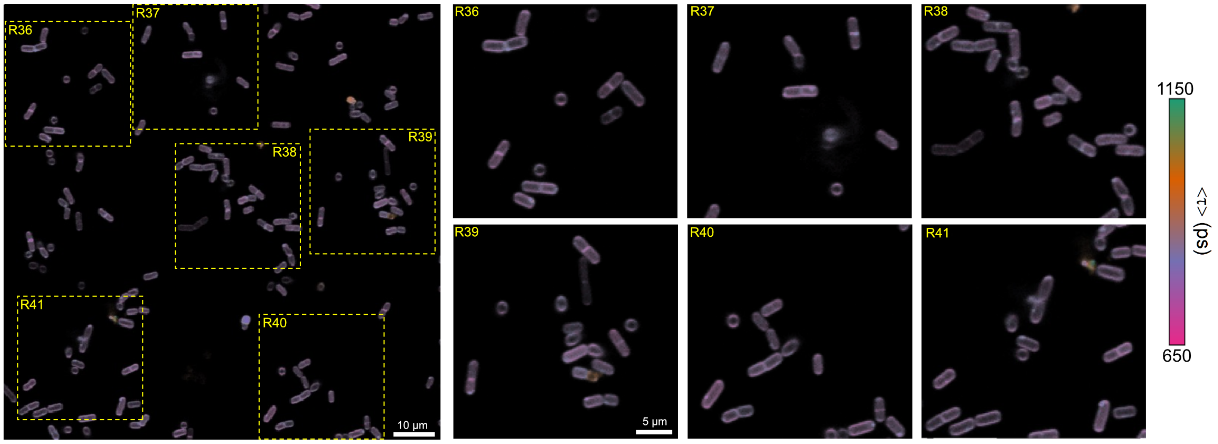

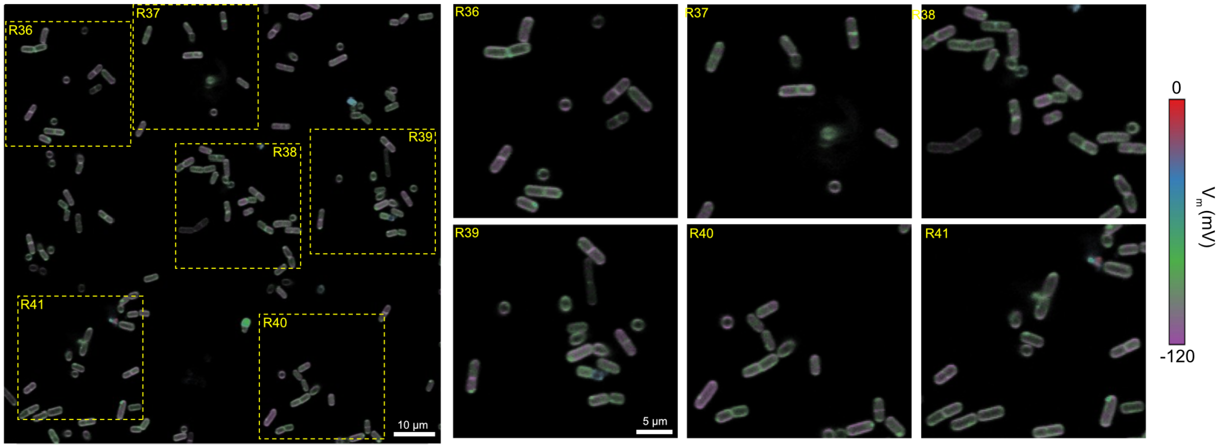

FOV8: unperturbed, in M9 only

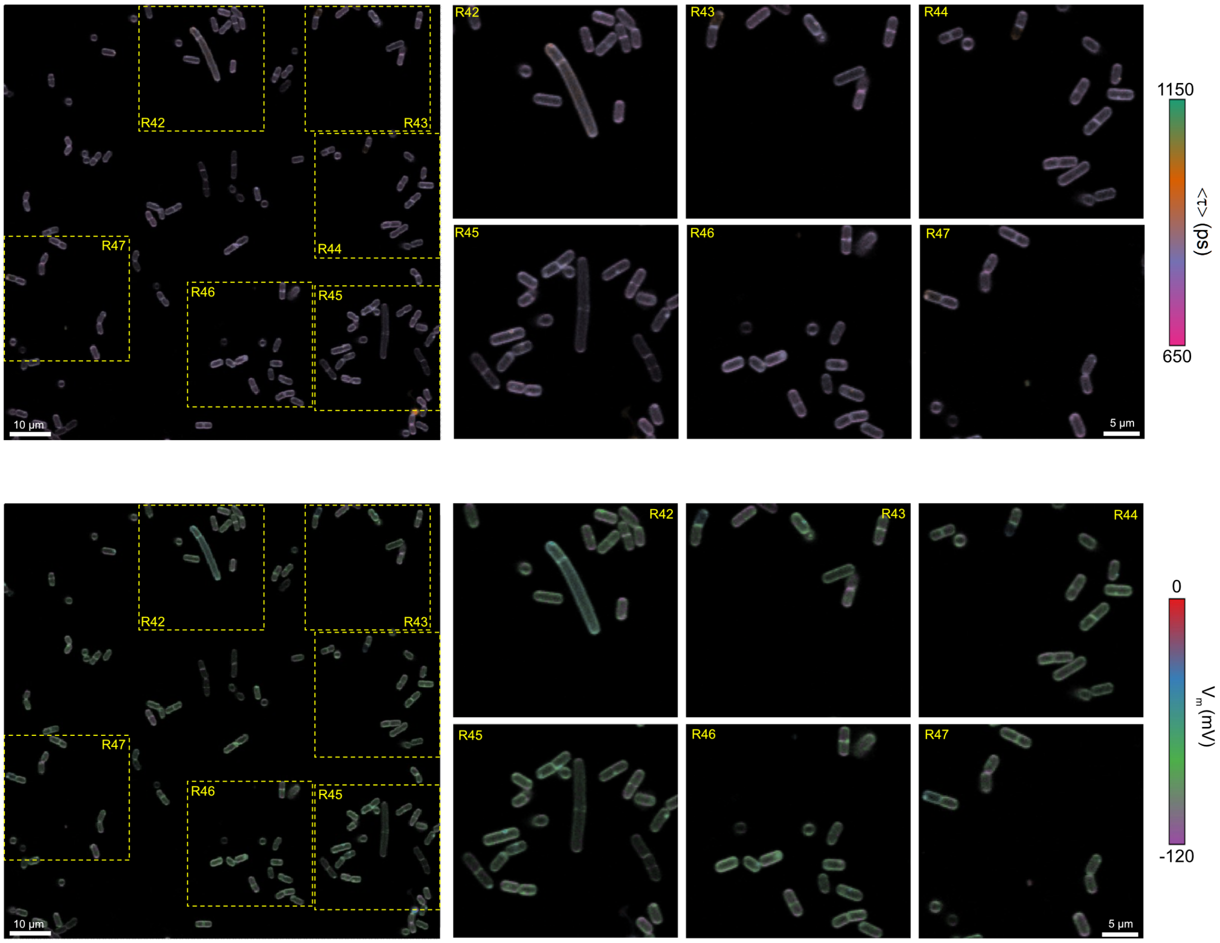

FOV9: unperturbed, in M9 only

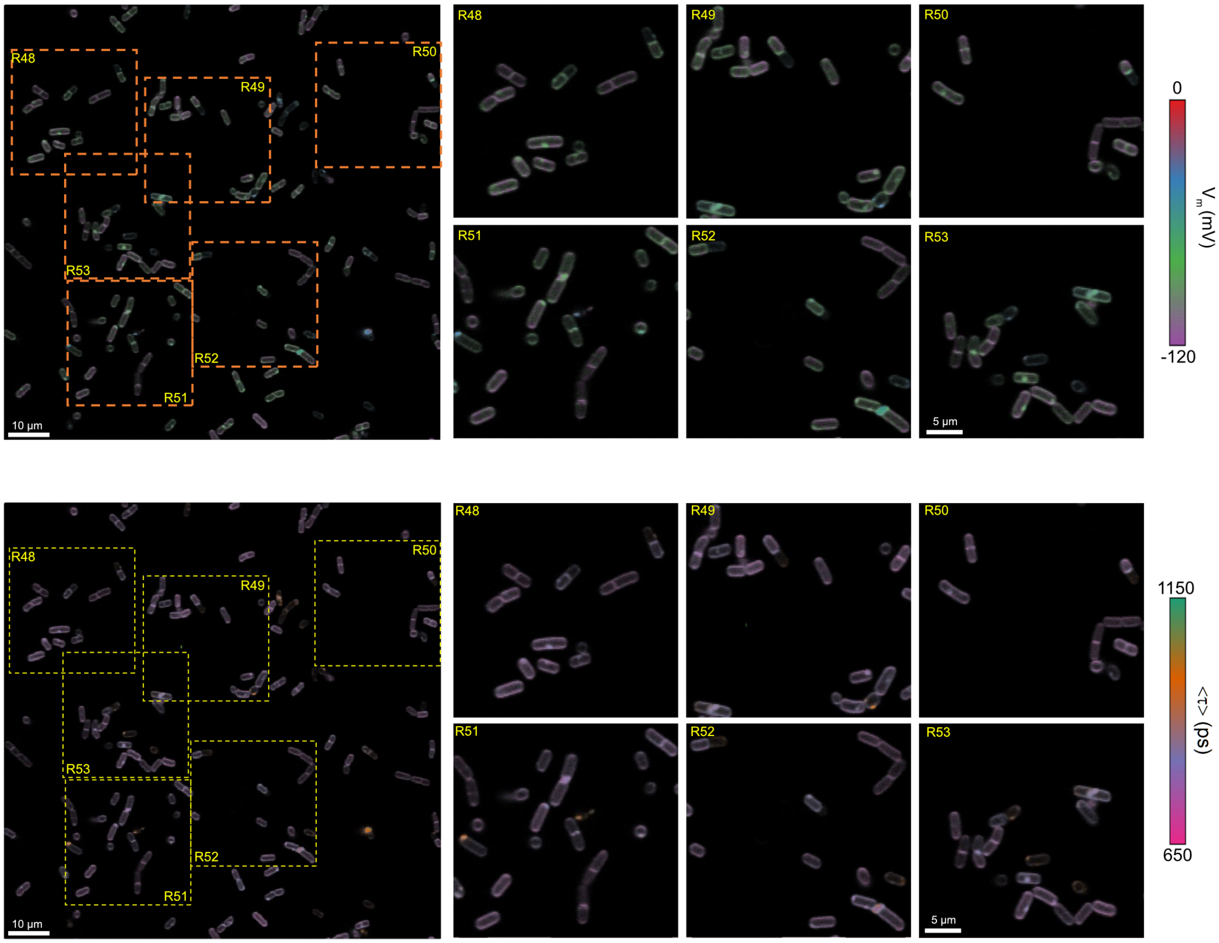

**Supplementary Figure 5**: Different field of views (FOVs) depicting the effect of varying extracellular chemical conditions on the average lifetime of VF2.1.Cl and membrane potential in the membrane of *B. subtilis*. (a) Unperturbed cells in MSgg only; FOV1-3: <τ> ~ 865 ± 24 ps; (b) Chemically depolarized cells in MSgg (25 μM valinomycin + 240 mM KCl); FOV4-6: <τ> ~ 1100 ± 46 ps; (c) Unperturbed cells in M9 only; FOV7-9: <τ> ~ 777 ± 14 ps; In each case, a phasor-based amplitude-averaged lifetime map is shown first, and the corresponding membrane potential map obtained by application of the calibration relation (Eq. S32) is shown next.

#### Supplementary Figure 6: Average lifetime dispersion across measurements, MSgg vs M9

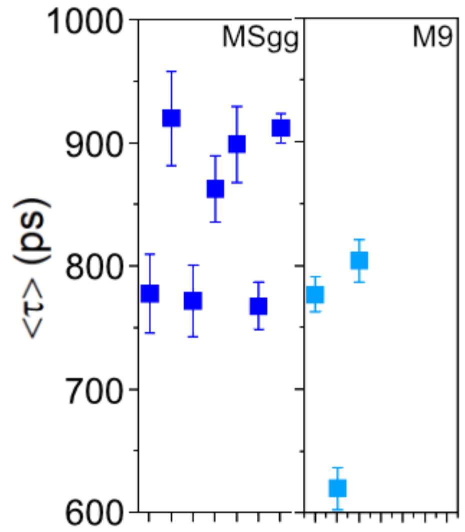

**Supplementary Figure 6**: Average lifetime dispersion across measurements and external chemical conditions.

### Supplementary Tables

#### Supplementary Table 1: MSgg composition

| Component | Concentration |
|---|---|
| Glycerol 0.5% (vol/vol) – C source | 68.4 mM |
| Monosodium glutamate 0.5% (wt/vol), monohydrate salt – N source | 29.5 mM |
| Thiamine-HCl | 2 µM |
| 3-(N-morpholino) propanesulfonic acid sodium salt, MOPS-Na pH 7.0 | 10 mM |
| $KH_2PO_4$ | 2.1 mM |
| $K_2HPO_4$ | 2.9 mM |
| $MgCl_2$, $6H_2O$ | 2 mM |
| $CaCl_2$, $2H_2O$ | 700 µM |
| $FeCl_3$, $6H_2O$ | 100 µM |
| $MnCl_2$, $4H_2O$ | 50 µM |
| $ZnCl_2$ | 1 µM |

Composition of minimal salts glycerol glutamate (MSgg) medium (1 L) after ref. (4)

#### Supplementary Table 2: VF2.1.Cl and VF2.0.Cl fluorescence decay fit parameters

Results of nonlinear least-square (weighted) fit (NLSF) of fluorescence decays in DMSO and *B. subtilis* membrane. The long and short lifetime components as well as fractional amplitudes are reported. Amplitude-weighted average lifetimes computed according to Eq. 7 in the main text are also indicated.

| Probe (medium) | NLSF lifetimes (in ns) and relative amplitudes | | |
|---|---|---|---|
| | $\tau_1$ ($a_1$) | $\tau_2$ ($a_2$) | $\tau_{avg}$ |
| VF2.0.Cl (DMSO) | 3.21 (1) | | 3.21 |
| VF2.0.Cl (membrane) | 2.72 (1) | | 2.72 |
| VF2.1.Cl (DMSO) | 2.90 (0.35) | 0.81 (0.65) | 1.53 |
| VF2.1.Cl (membrane) | 1.87 (0.27) | 0.32 (0.73) | 0.74 |

#### Supplementary Table 3: VF2.1.Cl and VF2.0Cl fluorescence decay fit parameters

Results of nonlinear least-square (weighted) fit (NLSF) of fluorescence decays of VF2.0Cl and VF2.1.Cl in various conditions in the absence of cells

| medium | Probe | NLSF lifetimes (in ns) and relative amplitudes | | |
|---|---|---|---|---|
| | | $\tau_1$ ($a_1$) | $\tau_2$ ($a_2$) | $\tau_{avg}$ |
| DMSO | VF2.1.Cl | 2.900 (0.35) | 0.811 (0.65) | 1.530 |
| | VF2.0.Cl | 3.210 | | 3.210 |
| MSgg | VF2.1.Cl | 2.950 (0.63) | 0.535 (0.37) | 2.057 |
| | VF2.0.Cl | 2.758 | | 2.758 |
| MSgg + Valinomycin | VF2.1.Cl | 3.040 (0.67) | 0.542 (0.33) | 2.200 |
| | VF2.0.Cl | 3.058 | | 3.058 |
| MSgg + Valinomycin + KCl | VF2.1.Cl | 2.770 (0.54) | 0.454 (0.46) | 1.693 |
| | VF2.0.Cl | 2.680 | | 2.680 |

#### Supplementary Table 4: Average lifetimes for VF2.1.Cl in cell under various extracellular chemical conditions (corresponding to Figure 6)

| Extracellular chemical condition | # | $<\tau>$ (ps) | $\sigma_\tau$ (ps) |
|---|---|---|---|
| MSgg media only | 1 | 778 | 32 |
| | 2 | 920 | 38 |
| | 3 | 772 | 29 |
| | 4 | 863 | 27 |
| MSgg + 25 µM valinomycin | 1 | 920 | 41 |
| | 2 | 822 | 54 |
| | 3 | 951 | 42 |
| | 4 | 925 | 39 |
| | 5 | 931 | 32 |
| | 6 | 899 | 30 |
| MSgg + 25 µM valinomycin + 240 mM KCl | 1 | 1129 | 44 |
| | 2 | 1107 | 43 |
| | 3 | 1126 | 40 |
| MSgg + 25 µM valinomycin + 300 mM KCl | 1 | 1075 | 38 |
| | 2 | 1210 | 43 |
| | 3 | 1029 | 70 |
| | 4 | 1066 | 54 |

#### Supplementary Table 5: Shot noise contribution to the total standard deviation of the average lifetime

| set | chemical condition | $<\tau>$ (ps) | $\sigma_{<\tau>}$ (ps) | $\sigma_{<\tau>}^{SN}$ (ps) | $\sigma_{<\tau>}^{Other}$ (ps) | $\sigma_{V_m}^{Other}$ (mV) |
|---|---|---|---|---|---|---|
| Series cal | MSgg+Val@25µM+KCl@1mM+NaCl@239mM | 786 | 51 | 20 | 47 | 38 |
| Series cal | MSgg+Val@25µM+KCl@10mM+NaCl@230mM | 848 | 48 | 18 | 45 | 9 |
| Series cal | MSgg+Val@25µM+KCl@40mM+NaCl@200mM | 872 | 46 | 25 | 39 | 8 |
| Series cal | MSgg+Val@25µM+KCl@80mM+NaCl@160mM | 1000 | 40 | 26 | 30 | 6 |
| Series cal | MSgg+Val@25µM+KCl@160mM+NaCl@80mM | 1081 | 41 | 21 | 35 | 7 |
| Series cal | MSgg+Val@25µM+KCl@240mM | 1129 | 44 | 23 | 38 | 8 |
| Series 1 | MSgg+Val@25µM+KCl@30mM+NaCl@270mM | 979 | 36 | 19 | 31 | 6 |
| Series 1 | MSgg+Val@25µM+KCl@150mM+NaCl@150mM | 961 | 44 | 20 | 39 | 8 |
| Series 2 | MSgg+Val@25µM+KCl@100mM+NaCl@200mM | 916 | 48 | 22 | 43 | 9 |
| Series 2 | MSgg+Val@25µM+KCl@60mM+NaCl@240mM | 900 | 58 | 20 | 54 | 11 |
| Series 2 | MSgg+Val@25µM+KCl@300mM | 1029 | 70 | 22 | 66 | 13 |
| Series 2 | MSgg+Val@25µM+KCl@30mM+NaCl@270mM | 772 | 60 | 17 | 58 | 46 |
| Series 4 | MSgg+Val@25µM | 931 | 32 | 27 | 17 | 4 |
| Series 4 | MSgg+Val@25µM+KCl@30mM | 1010 | 45 | 22 | 39 | 8 |
| Series 5 | MSgg+Val@25µM+KCl@240mM | 1107 | 43 | 28 | 33 | 7 |
| Series 5 | MSgg+Val@25µM+KCl@150mM | 1088 | 40 | 33 | 23 | 5 |

The last 5 columns of the table report the population-mean amplitude-averaged lifetime obtained by phasor analysis, as well as the corresponding standard deviation. $\sigma_{\langle\tau\rangle}^{SN}$ is the simulated shot noise limited standard deviation of the amplitude-average lifetime (for cells with total intensity corresponding to the mode of the population intensity histogram). $\sigma_{\langle\tau\rangle}^{Other}$, given by Eq. (5) in the main text, corresponds to the standard deviation contribution due to sources other than shot noise, presumably physiological state differences between cells. $\sigma_{Vm}^{Other}$ converts the latter into mV using the calibration curve of Figure 4c.

#### Supplementary Table 6: Data for the discussion of Figure 7

| Change | $\Delta[K^+]_{out}$ (mM) | $\Delta$ [Xyl] (mM) | $\Delta Osm_{out}$ (mV) | $\Delta[K^+]_{out}$ | $\Delta[K^+]_{in}$ | Predicted $\Delta V_m$ | Observed $\Delta V_m$ (mV) |
|---|---|---|---|---|---|---|---|
| 2→3 | -60 | +120 | 0 | - | 0 | - | -91 |
| 5→6 | -60 | +120 | 0 | - | 0 | - | +47 |
| 1→3 | 0 | +120 | +120 | 0 | + | - | -75 |
| 3→6 | 0 | +180 | +180 | 0 | + | - | +71 |
| 1→6 | 0 | +300 | +300 | 0 | + | - | -4 |
| 2→4 | 0 | +120 | +120 | 0 | + | - | +12 |
| 4→5 | 0 | +60 | +60 | 0 | + | - | -79 |
| 2→5 | 0 | +180 | +180 | 0 | + | - | -67 |
| 7→8 | 0 | +100 | +100 | 0 | + | - | -30 |
| 3→4 | +60 | 0 | +120 | + | + | | +103 |
| 5→7 | +40 | +20 | +100 | + | + | | +72 |

Parameters corresponding to the different virtual transitions (i → j) from one external condition to another. The condition index (i or j) correspond to the list of Table 1 in the text. The change in $[K^+]_{out}$, [Xyl] and medium osmolality are indicated in the 2nd, 3rd and 4th columns. Qualitative changes for $[K^+]_{out}$,and $[K^+]_{in}$ are indicated in the next columns (+: increase, -: decrease, 0: no change), as well as the qualitative change in $V_m$ predicted by a simple argument discussed in the main text. The last column reports the measured $\Delta V_m$ and indicates by a color code whether it matches the qualitative prediction (green) or not (red), or is compatible with either within the experimental uncertainty (orange). The last two transitions are not colored due to the impossibility to predict the membrane potential change based on our simple argument.

### Supplementary Notes

#### Supplementary Note 1: *B. subtilis* culture, growth and membrane staining protocols

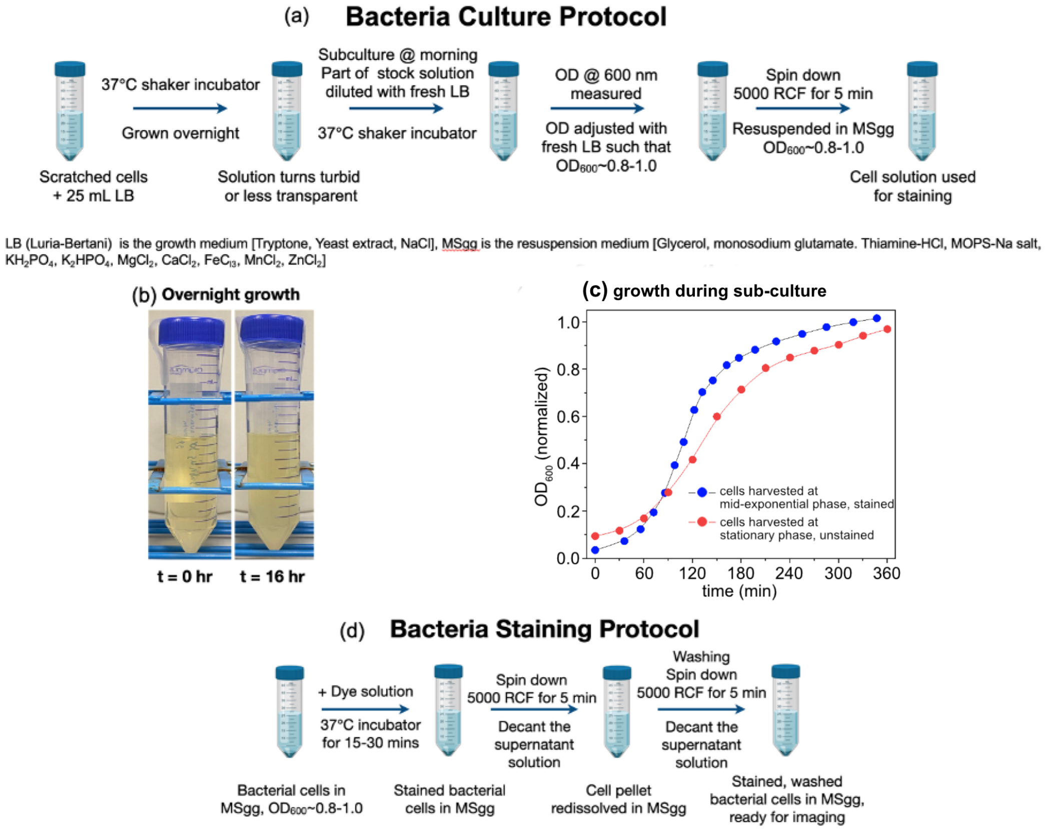

**Supplementary Figure 7**: Overview of sample preparation steps.

The frozen stock of cells was grown overnight in 25 mL of LB medium in a shaking incubator at 37°C. 1 mL aliquot of overnight grown cells was withdrawn, added to 25 mL of fresh LB medium and grown to mid-log phase ($OD_{600}$ = 0. – 1.0). The OD was measured with PerkinElmer instruments Lambda 25 UV/VIS spectrometer. The growth curve (during subculture) for cells harvested at stationary phase depicted in Supplementary Figure 6c (red). 1 mL of bacterial cells at the mid-exponential phase were incubated (for ~30 min) with voltage probe VF2.1.Cl (bath concentration ~100 nM). The cell's ability to grow and divide after staining was tested. 1 mL of stained cells resuspended in 25 mL of fresh LB medium in a shaking incubator at 37°C. The stained cells continue to grow and divide. The growth curve (during subculture) for cells harvested at mid-exponential phase, stained with VF2.1.Cl, is depicted in Supplementary Figure 6c (blue). Staining with VF2.1.Cl did not exhibit any growth-inhibitory effects. Stained bacterial cells were resuspended in MSgg media for imaging. Cells were adhered to poly-L-lysine treated Petri dish. Cells were then allowed 30 min to settle down on the glass surface.

In a given day when a series of experiments were performed for different extracellular chemical conditions (such as varying concentration of KCl and Xylose), samples were imaged randomly to try and eliminate the effect of time-elapsed since initial preparation until observation.

#### Supplementary Note 2: Imaging details

#### FLIM data acquisition details

FLIM datasets were acquired using a Leica SP8 LIGHTNING confocal laser scanning microscope (Advanced Light Microscopy and Spectroscopy - ALMS - Core Facility at the Center for NanoSystems Institute - CNSI, UCLA) (RRID:SCR_022789). An oil immersion, high-NA objective (Leica HC PL APO 63x/1.40 OIL and 100x/1.40 OIL) was used. Excitation was provided by a tunable white light supercontinuum pulsed laser (WLC 470-670 nm) at 78 MHz (<100 ps FWHM). An acousto-optic beam splitter (AOBS) and HyD SMD hybrid detector were used to collect emitted photons. For our experiments with VoltageFluors, $\lambda_{ex}$ was set to 514 nm and the detection range was set to 536-700 nm. A high precision motorized stage was used to move the sample around. A resonant galvo-mirror was used to acquire raster-scanned 512×512 pixel datasets with a 3.16-7.69 µs dwell time per pixel. To increase signal, 15-25 images were accumulated for each field of view. An OkoLab stage-top enclosure maintained a 37 °C temperature during live cell imaging. Raw TCSPC data was exported in the time-tagged time-resolved ptu format, converted using AlliGator into accumulated series of time-binned images with the native resolution of 97.1 ps, and exported into the AlliGator HDF5 FLI dataset open file format(5). A reference dataset was obtained using a dilute solution of Allura Red in water imaged in the same conditions as the sample.(6) Due to its very short lifetime (~10 ps) and broad absorption spectrum, it is a convenient sample to obtain an approximate instrument response function (IRF) including the optical characteristics of the setup (laser pulse width, optics dispersion) as well as detector and electronics responses.

#### White light image acquisition details

White light images were acquired using a Nikon widefield microscope (Nikon Eclipse Ti, having a 1.5x magnification changer) equipped with an oil immersion, high-NA objective (Nikon Plan Apo VC 60x/1.40 oil). Illumination was provided by a 100 W halogen lamp for white light and a 500 mW Lumencor VCGR-01B PS Aura 4-Color Light Engine for fluorescence imaging. An Andor iXon+ 897 EMCCD camera (pixel size 16µm) was used to capture images with an exposure time of 100 ms. Post-processing, including contrast adjustment, was performed using the free software Fiji.

White light images were also acquired using a Leica SP8 Confocal Microscope, switched to white light mode through Leica LAS X software. An oil immersion, high-NA objective (Leica HC PL APO 63x) was used. The transmitted light source provided white light illumination. A camera was used to capture images with an exposure time of 10 ms. Post-processing, including contrast adjustment, was performed using Fiji.

#### Supplementary Note 3: Effect of imaging conditions/light exposure on B. subtilis cell growth (viability after imaging)

In the imaging experiments/FLIM measurements we describe in the manuscript, cells within a given FOV remained typically exposed to green pulsed laser light ($\lambda_{ex}$ – 514 nm) for 1 to 2 min depending on the desired signal. In order to test stained cell's ability to grow under single-shot and/or continuous light exposure we performed time-lapse imaging of a given FOV under (a) widefield CW illumination and (b) confocal pulsed illumination separately. VF2.1.Cl stained bacterial cells were resuspended in LB (conducive to growth) in a flow channel.

We first exposed cells to wide-field white light illumination and performed time-lapse observation of their growth. Under widefield illumination, the entire FOV was exposed to light simultaneously. In the middle of this recording (at the $55^{th}$ minute), we exposed the entire FOV of cells to a green LED for 2 min and resumed the time-lapse observation under white light illumination. The entire FOV was exposed to green light passed through a YFP filter cube (excitation filter 500/24, dichroic 520, emission filter 542/27). Time lapse (~15 min interval) images were acquired for ~2hr to monitor the growth, as depicted in Supplementary Figure 8. At the 55th minute, the whole FOV is exposed to light irradiation for 2 min. Cells continue to grow after the 57th minute as can be seen from the images in Supplementary Figure 8. While this imaging modality is not used in the manuscript, it is provided for information, as it is a common modality and easier to perform for most laboratories although it only allows performing quantum yield change measurements, not lifetime measurements as is the focus of our work.

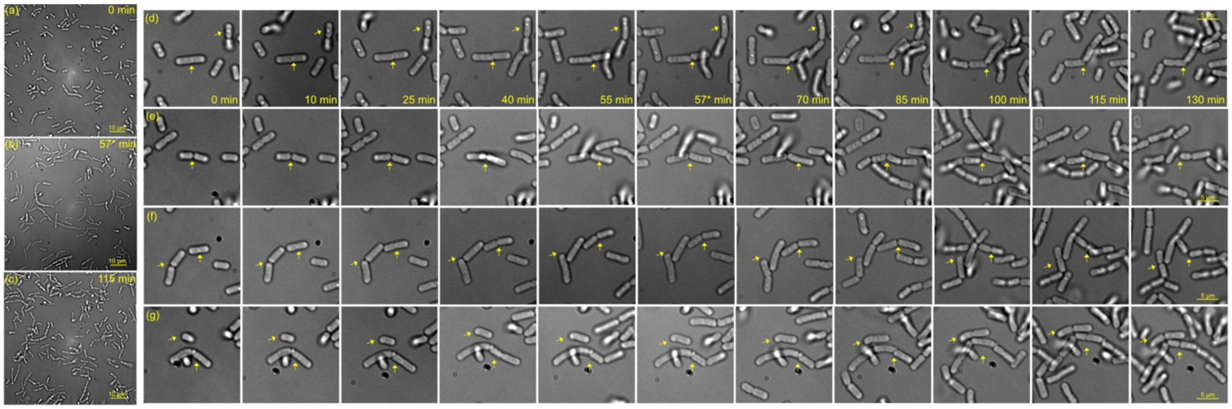

**Supplementary Figure 8**: (a)-(c) Full FOV BF images of VF2.1Cl stained B. subtilis cells resuspend in LB medium. Scale bar 10 µm. The FOV (89 µm x 89 µm) exposed to light irradiation (through YFP filter) for 2 minutes at the 55[th] minute. Cell density continue to increase over time even after light exposure. (d)-(g) Zoomed region of interest (22 µm x 22 µm) within the same FOV. Scale bar 5 µm. Individual cells are observed to grow and divide over time. Some cells of interest indicated with arrow for ease of viewing.

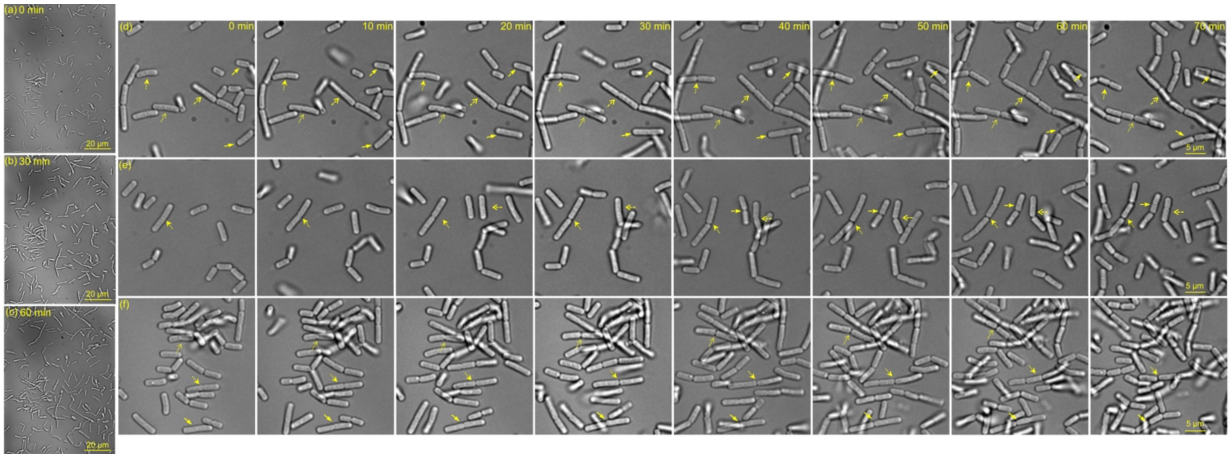

**Supplementary Figure 9**: (a)-(c) Full FOV BF images of VF2.1Cl stained B. subtilis cells resuspend in LB medium. Scale bar 20 µm. The FOV undergoes repeated (10 min interval) exposure to laser irradiation for ~1-2 mins. Cell density continue to increase over time even after light exposure. (d)-(f) Zoomed region of interest within the same FOV. Scale bar 5 µm. Individual cells are observed to grow and divide over time. Some cells of interest indicated with arrow for ease of viewing.

In a separate experiment, cells were exposed to pulsed confocal illumination. Cells in LB medium were exposed to raster-scanning confocal excitation with a 514 nm pulsed laser beam (~ 100 ps FWHM, 78 MHz). Scans were comprised of 512 x 512 pixels, with ~7 µs/pixel dwell time. In order to improve signal-to-noise ratio, we acquired and summed multiple frames (20), for a total duration of ~1-2 min. However, it is worth noting that each pixel remained exposed to light for <150 µs. Time-lapse confocal images were acquired every ~10 min for ~1hr to monitor the growth, together with white light images. Representative images are shown in Supplementary Figure 9 (white light) and Supplementary Figure 10 (confocal). Unlike in our wide-field illumination experiments, each FOV was exposed to 514 nm pulsed laser light every ~10 min, a much harsher treatment than in the previously described measurements. However, as can be seen from the images in Supplementary Figure 9-10, cells continued to grow after repeated laser irradiation. Note that FLIM measurements were performed in a minimal medium (MSgg, M9) not conducive to growth.

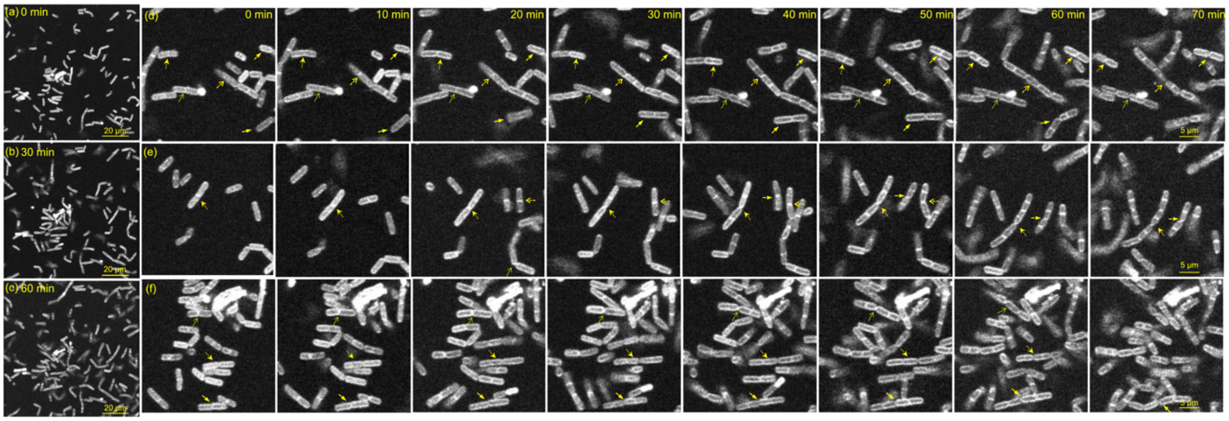

**Supplementary Figure 10**: (a)-(c) Full FOV confocal images of VF2.1Cl stained B. subtilis cells resuspend in LB medium. Scale bar 20 μm. The FOV undergoes repeated (10 min interval) exposure to laser irradiation for ~1-2 mins. Cell density continue to increase over time even after light exposure. (d)-(f) Zoomed region of interest within the same FOV. Scale bar 5 μm. Individual cells are observed to grow and divide over time. Some cells of interest indicated with arrow for ease of viewing.

#### Supplementary Note 4: Additional details for phasor analysis

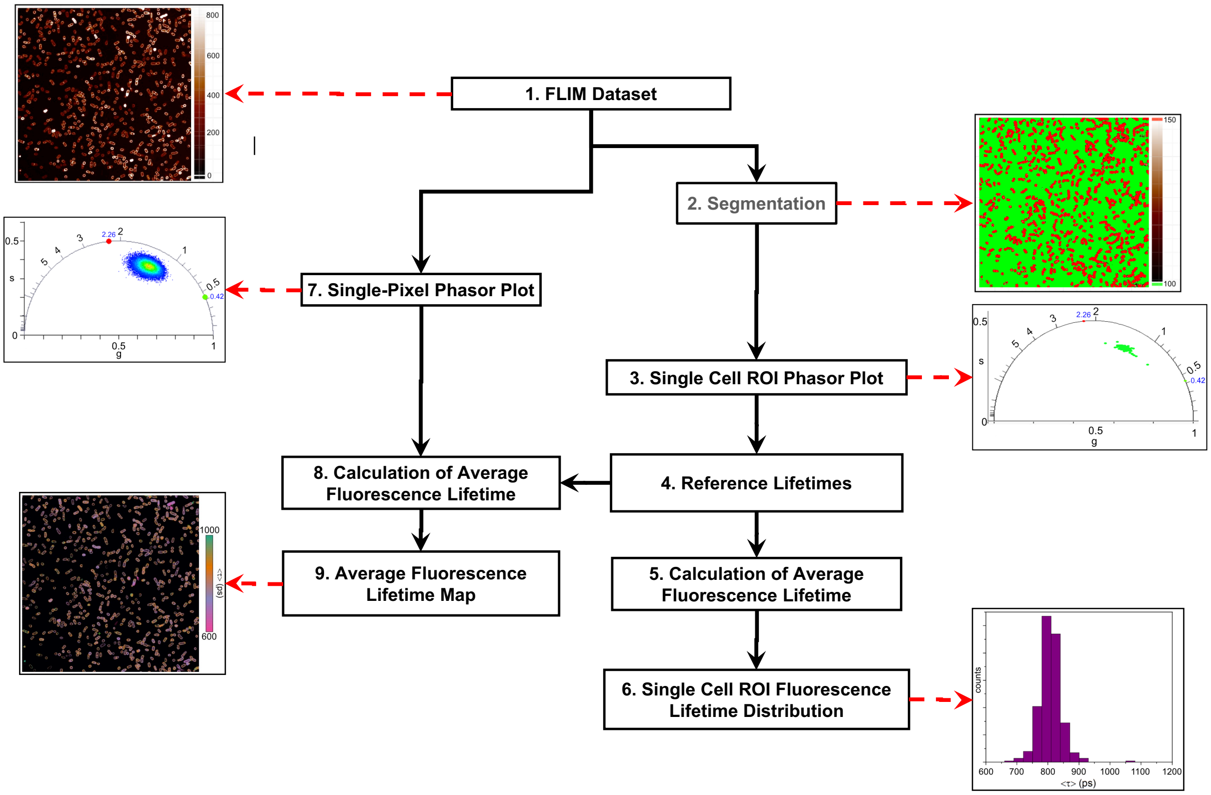

**Supplementary Figure 11**: overview of the different analysis steps involved in our workflow.

Phasor analysis, a faster and more robust approach introduced by Gratton and others, provides a simple yet powerful way to visualize and quantify the relevant information without fitting and is therefore ideally suited for handling the large amount of data encountered in FLIM (512 × 512 fluorescence decays/image).

**1. Dataset conversion**: Leica FLIM datasets were exported in the time-tagged ptu file format in the LASX software and converted into series of binned (or gate) images corresponding to photons arriving within a particular bin of 97.1 ps after the laser pulse in AlliGator. The corresponding datasets were saved individually in the AlliGator FLI dataset file format based on the open HDF5 format (see AlliGator's manual for details).(5)

**2. Bacterial cell segmentation**: Approximate cellular contours were obtained using the StarDist plugin(7) in the open source software Fiji.(8) In brief, the intensity (sum of all gates) image's contrast was adjusted to the min and max 1-percentile and exported as a 16-bit TIFF image. The StarDist algorithm default parameters were used to identify ROIs (see Supplementary Figure 8). The corresponding ROI image was then saved and reimported in AlliGator as a mask image to define the corresponding ROIs.

**3. Single-Cell Phasor Analysis**: For each cell ROI, the total fluorescence decay of all encompassed pixels was computed. As discussed in Supplementary Note 6, the signal collected in pixels located inside the cell most likely originates from the membrane above and below the focal plane and therefore correspond to the signal from VF dyes exposed to the cell's membrane potential. Phasors $z$ corresponding to these fluorescence decays were computed and calibrated with the IRF's phasor $z_{ref}$ (Allura Red decay,(6) identified with lifetime 0 and calibrated phasor $z_0 = 1$) according to:

$$z_{cal} = z / z_{ref} \tag{S1}$$

using the Analysis→FLI Dataset→Multiple ROIs Analysis→All ROIs Phasor Analysis→Non-Interactive (Fast) AlliGator script.

**4. Reference Lifetimes Definition**: The calibrated phasors of cells exposed to a given condition were analyzed globally to define "natural" single-exponential references for this condition. Briefly, considering the elongated (elliptical) distribution of observed phasors, we assumed that the phasors were representative of a distribution of bi-exponential decays characterized by the same single-exponential decay components (with lifetimes $\tau_1$ and $\tau_2$), but different amplitudes. Since such phasors are expected to be located on the segment connecting the two single-exponential decay phasors $z_{j,\, j=1 \text{ or } 2} = \left(1 - i2\pi f \tau_j\right)^{-1}$, a "natural" definition for this segment is that of the ellipse's major axis. Single lifetime phasor references $z_1$ and $z_2$ were defined as the intersections of this major axis with the universal semicircle (UC), locus of single exponential decay phasors, defined by $\left(g - 1/2\right)^2 + s^2 = \left(1/2\right)^2$ (AlliGator's Phasor Graph Phasor Ratio Reference→Use Selected Phasor Plots Major Axis/UC Intersections function).

**5. Calculation of Single-Cell Amplitude-Averaged Fluorescence Lifetime**: For each single-cell phasor, the intensity fraction $f_1$ of reference 1 was then given by:(9)

$$f_1 = \left(z_{cal} - z_1\right) \frac{\left(z_1 - z_2\right)^*}{\left|z_1 - z_2\right|^2} \tag{S2}$$

where $z^*$ indicates the complex conjugate of $z$. This expression corresponds to the fractional distance of the orthogonal projection of the calibrated phasor $z_{cal}$ onto the segment connecting $z_1$ and $z_2$.
The amplitude fraction needed to computed the amplitude-averaged lifetime $<\tau>_a$ was then obtained by the conversion formula: (9)

$$\langle \tau \rangle_a = \left(f_1/\tau_1 + f_2/\tau_2\right)^{-1} \tag{S3}$$

**6. Single-Cell Fluorescence Lifetime Distribution**: The corresponding distribution was histogrammed and its mean and standard deviation computed (AlliGator Phasor Graph's Parameter 2 vs Parameter 1 Scatter Plot function followed by AlliGator's Lifetime & Other Parameters Graph Plot Histogram function).

**7. Single-pixel Phasor Analysis**: To compute the pixel-level phasor plot (used in the next analysis step), single-pixel decays limited to pixels included in the cell ROIs were converted to calibrated phasors using Eq. (S1) and represented as a 2-dimensional ($g$, $s$) color-coded histogram (AlliGator's Phasor Plot with Limit Phasor Plot Calculation to Selected Image ROI(s) option).

**8. Single-pixel Amplitude-Average Lifetime**: Using the single-pixel phasors calculated in the previous step (7), and the phasor references defined in step 4, individual pixel amplitude-averaged lifetime were computed using Eq. (S3).

**9. Color-coded Average Lifetime Map**: Using a 256 color-scale mapped to a lifetime range [$\tau_{min}$, $\tau_{max}$], the cell ROI pixels were colored accordingto their computed lifetime, the brightness of each pixel being proportional to the pixel intensity (after user-defined contrast adjustment).

**Single-Cell NLSF Analysis and Amplitude-Average Lifetime Calculation**: The decays of single-cell ROIs were fitted with a two-exponential model after convolution with the IRF as defined in Eq. (6) of the main text, using the Analysis→FLI Dataset→Multiple ROIs Analysis→All ROIs NLSF Analysis→Non-Interactive (Fast) AlliGator script. The guess parameters used to initialize the fit were computed heuristically based on the actual decay. No parameter was constrained. The corresponding amplitude-averaged lifetime defined by Eq. (7) of the main text was histogrammed and the resulting histogram fitted with a normal distribution $N(\bar{\tau},\sigma_{\tau})$.

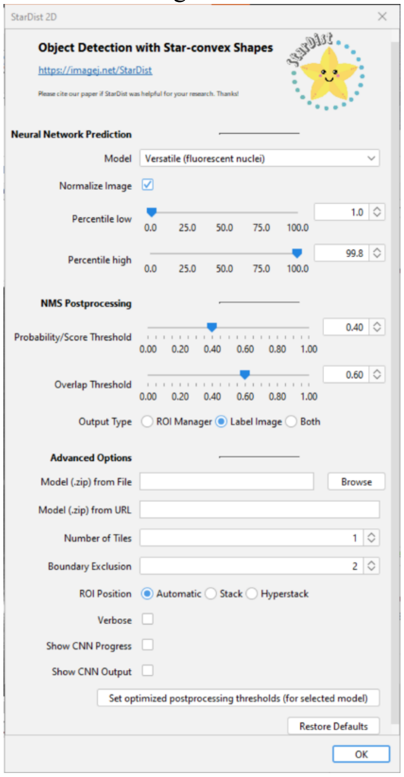

**Supplementary Figure 8**: ImageJ StarDist plugin (default) parameters used in this work.

### Supplementary Note 5: Shot noise influence on phasor-based amplitude-averaged lifetime

#### Principle of the estimation

To assess the contribution of shot noise to the uncertainty on the amplitude-averaged lifetime computed in step 5 of Supplementary Note 4, we processed each single cell fluorescence decay in the following manner.

A single-cell decay is defined by $G$ arrival time bin values $\{I(t_i)\}_{i=1,\ldots,G}$, each being ideally a random variable drawn from a Poisson distribution with mean value $S_T(t_i)$, where $S_T(t)$ is defined by Eq. (6) in the main text. In the absence of this randomness, fitting $S_T(t)$ by a bi-exponential decay model (convolved with the IRF) should return the exact value of the decay parameters (assuming a numerically efficient NLSF algorithm). Because of the randomness associated with the finite value of each bin $I(t_i)$ and their statistical nature, the fitted parameters are generally different from the true ones. Repeating the measurement $M$ times would provide an estimate of the dispersion of each parameter $P_j$ (ideally normally distributed with mean value $\bar{P}_j$ and standard deviation $\sigma_{P_j}$). Instead, we simulated $M$ = 1,000 replicas of the single measured decay, replacing each $I(t_i)$ by a random number Poi($I(t_i)$) drawn from the Poisson distribution with average value $I(t_i)$, and computed the corresponding <τ> based on the phasor references defined in step 4 of Supplementary Note 4. The corresponding standard deviation $\sigma_{<\tau>}^{SN}$ of the distribution of amplitude-averaged lifetime was used as an estimate of the uncertainty on <τ> due to shot noise, computed for each single-cell decay.

Note that this is not exactly equivalent to repeating a (simulated) measurement, where the Poisson distribution mean value would be $S_T(t_i)$ instead of $I(t_i)$, which would require fitting the decay with the model $S_T(t_i)$. To avoid fitting the decay prior to this analysis (which would in any case not guarantee that the true value $S_T(t_i)$ is obtained), we used $I(t_i)$ instead. To verify that this replacement led to a reasonable estimation of the shot noise influence, we compared the amplitude-averaged lifetime distributions obtained in both cases, assuming a known underlying bi-exponential decay and total number of photons $S$, repeating the calculation $N$ times (number of simulations), using the AlliGator Analysis→Tools→ Shot Noise Influence on Average Lifetime tool (Supplementary Figure 12 & 13).

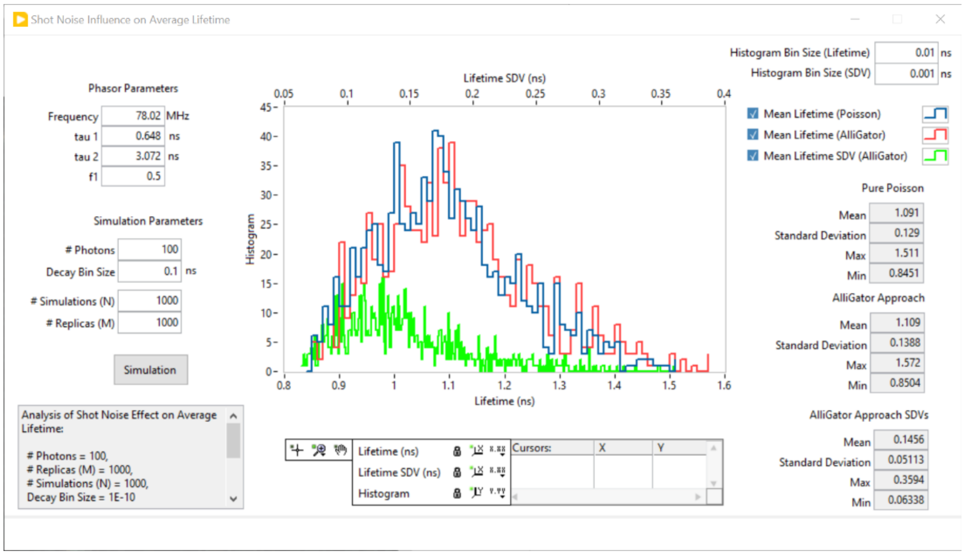

**Supplementary Figure 12:** Screenshot of the Shot Noise Influence on Average Lifetime tool in AlliGator. The simulation parameters: # Replicas (M) and # Simulations (N) correspond to those discussed in the text. The user can specify the type of decay to study by providing single-exponential references τ1 and τ2, intensity fraction of the first reference f1 and phasor frequency f, as well as the total number of photons in the simulated decays (# Photons). To reproduce the type of binned TCSPC data used in this study, the Decay Bin Size needs to be provided as well. The output of the simulation consists of 3 curves and associated statistics: 1) Mean Lifetime (Poisson) is the distribution of average lifetimes obtained from simulating N decays with the specified number of photons;, 2) Mean Lifetime (AlliGator) is the distribution of average lifetime obtained from processing the previous N simulated decays as described in the text, namely creating M replicas of each and computing the mean lifetime and standard deviation of the results; 3) Mean Lifetime SDV (AlliGator) is the distribution of lifetime standard deviations obtained in the previous process. The particular example shown here corresponds to one of the measured lifetimes in the main text. # Photons = 100 corresponds to a number of photon representative of a single-pixels signal. The mean standard deviation of ~150 ps (upper horizontal scale) is fairly large compared to the actual lifetime of 1.1 ns.

Experiments with various combinations of parameters indicate that there is no significant difference between the two estimates, justifying the use of our simplified approach to compute the influence of shot noise on the computed amplitude-average lifetime (see caption of Supplementary Figure 12).

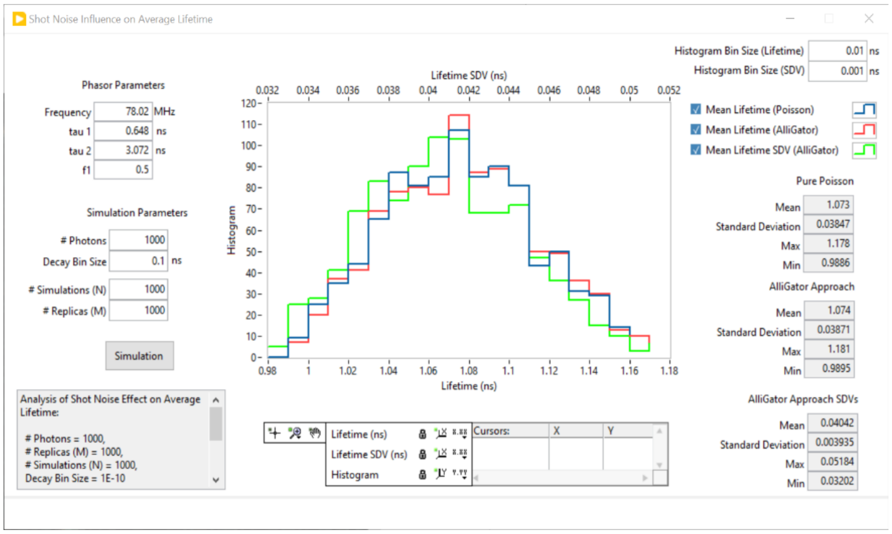

**Supplementary Figure 13:** Screenshot of the Shot Noise Influence on Average Lifetime tool in AlliGator. The parameters are identical to those in Supplementary Figure 12 except for the number of photons (1000 instead of 100). The observed standard deviation (40 ps) is ~10-1/2 that observed in Supplementary Figure 12, as expected. This number of photons correspond to one of the dimmest cells typically observed in the experiments described in the main text.

#### Shot noise contribution to the observed average lifetime dispersion

To understand the relative contribution of shot noise to the observed distribution of lifetimes within a cell population submitted to a given set of conditions, we assumed that it acted as an independent random variable and that it, as well as the residual contributions, resulted in a normal distribution of lifetimes. In these conditions, the two normal distributions need to be convolved with one another and the resulting normal distribution has a variance equal to the sum of the independent variances (that due to shot noise and that due to all other residual contributions):

$$\sigma_{\langle\tau\rangle}^{2} = \left(\sigma_{\langle\tau\rangle}^{Other}\right)^{2} + \left(\sigma_{\langle\tau\rangle}^{SN}\right)^{2} \quad \text{(S4)}$$

The residual standard deviation, $\sigma_{\langle\tau\rangle}^{Other}$, is then given by Eq. (5) in the main text:

$$\sigma_{\langle\tau\rangle}^{Other} = \sqrt{\left(\sigma_{\langle\tau\rangle}\right)^{2} - \left(\sigma_{\langle\tau\rangle}^{SN}\right)^{2}} \quad \text{(S5)}$$

Because our experiments involved measurements of cells with significantly different intensities, and because $\sigma_{\langle\tau\rangle}^{SN}$ depends on intensity (being approximately proportional to $N^{-1/2}$, where $N$ is the total decay signal), Eq. (S5) needs to be evaluated for each cell individually, or alternatively, for groups of cells with similar signal. The latter analysis (called Compute Sliced Mean & SDV Plots in AlliGator) was used to generate the plots in Supplementary Figure 14a-c, which represent 3 examples (of cells in different conditions) of the observed total average lifetime standard deviation, $\sigma_{\langle\tau\rangle}$, and the shot noise contribution, $\sigma_{\langle\tau\rangle}^{SN}$, as a function of intensity, using 1000-photon wide intensity windows. As expected, the shot noise contribution decays as the square root of the intensity, while the total standard deviation obeys a much more random relation, sometimes decreasing as the shot noise contribution, but sometimes increasing, depending on the measurement. Calculating the difference (Eq, (S5)), it is possible to represent these data synthetically by computing a weighted average of the residual standard deviation as:

$$\left\langle \sigma_{\langle\tau\rangle}^{Other} \right\rangle = \frac{\sum_{i=1}^{p} q_i \sigma_{\langle\tau\rangle}^{Other}(S_i)}{\sum_{i=1}^{p} q_i} \tag{S6}$$

where $S_i$ is the average intensity in intensity window $i$, and $q_i$ the number of cells whose intensity falls within the bounds of window $i$ (the $q_i$'s are represented as gray histograms in Supplementary Figure 14a-c). This number is representative of the typical average lifetime dispersion due to all effects distinct from shot noise, which we tentatively attribute to cell-to-cell variability. A similar weighted average can be computed for $\sigma_{\langle\tau\rangle}$, both being represented relative to one another in Supplementary Figure 14d.

It is clear from this analysis that, while shot noise contributes to the observed average lifetime distribution (and in a few cases, such as shown in Supplementary Figure 14a, appears to account for all the observed dispersion), a significant dispersion is caused by other sources of variations.

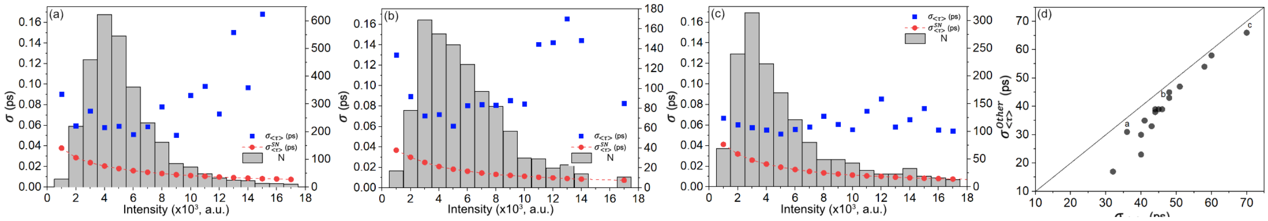

**Supplementary Figure 14**: (a-c) Examples of intensity-sliced analysis of the total lifetime standard deviation, $\sigma_{\langle\tau\rangle}$ (black squares) and the shot noise contribution to this standard deviation, $\sigma_{\langle\tau\rangle}^{SN}$ (red dots). The 3 plots correspond to representative datasets indicated by (a-c) in graph d. The gray histograms represent the number of cells in each intensity-slice (each slice is 1000-photon wide), read on the right axis of each graph. (d) Representation of the weighted-average residual standard deviation $\left\langle \sigma_{\langle\tau\rangle}^{Other} \right\rangle$ (Eq. (S6)) *vs* the equivalent quantity computed with the total standard deviation. The diagonal indicates identity between the two quantities.

#### Supplementary Note 6: 3-state model of the photophysics of VoltageFluor dyes

As outlined in the text, the fluorescence decays for the VF2.0.Cl and VF2.1.Cl dyes can be described by a 2- and 3-state model, respectively, analogous to that discussed by Li.(10) A simplified Jablonski diagram is provided in Supplementary Figure 15.

Fluorescence in VF2.0.Cl involves radiative and non-radiative de-excitation pathways from the first excited state 1 to the ground state 0, with a total rate $k_{10}$, equal to the inverse of the fluorescence lifetime $\tau_{2.0}$ of VF2.0.Cl, characterized by a single-exponential decay.

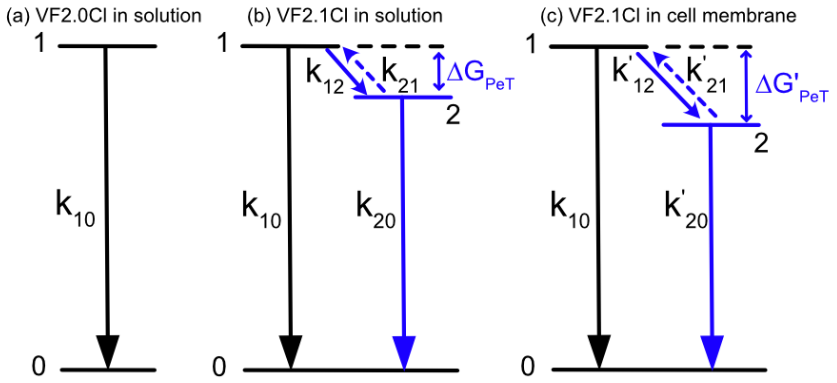

**Supplementary Figure 15**: Simplified Jablonski diagrams for (a) VF2.0.Cl in DMSO (b) VF2.1.Cl in DMSO, and (c) VF2.1.Cl in bacterial membranes. The PeT rate constant $k_{12}$ can be described by the *Marcus-Levich-Jortner's* formalism (Eq. (S7)). $\Delta G_{PeT}$ is the free energy change during PeT in the absence of external electric field. $\Delta G'_{PeT}$ is the external electric field E induced free energy change during PeT.

The photophysics of VF2.1.Cl involves a similar local excited state 1 (LE) for the donor-acceptor pair, akin to the 1 state of VF2.0.Cl, but also an additional intermediate state 2 for the charge-separated donor-acceptor pair, whose energy depends on the local electric field $\vec{E}$ . The availability of two separate relaxation channels from the LE state results in a bi-exponential decay, with a shorter average fluorescence lifetime and lower fluorescence quantum yield for VF2.1.Cl in the presence of an electric field. The dependency of $k_{12}$ (decay rate from 1 to 2) and $k_{20}$ (decay rate from 2 to 0) on the electric field (see below) results in a dependence of the fluorescence decay on $V_m = E/d$, where $d$ is the membrane thickness, the electric field $E$ being assumed uniform within and perpendicular to the membrane.

In the 3-state model described by Li (10) and schematically illustrated in Supplementary Figure 15, the PeT rate constant $k_{PeT} = k_{12}$ for the $1 \to 2$ transition can be formally computed using the Marcus-Levich-Jortner's formalism:(10-12)

$$k_{12} = A\sum_{n=0}^{\infty}\frac{S_c^n}{n!}\exp\left[-\frac{\left(\Delta G_{PeT}+\lambda_m+nh\omega_c\right)^2}{4\lambda_m k_B T}\right], \tag{S7}$$

where $A$ is a species-dependent constant depending on the electronic coupling constant between state 1 and 2, $\lambda_m$ is the reorganization energy of the surrounding medium during electron transfer, $\omega_c$ the average of the high-frequency intramolecular vibrational modes facilitating the electron transfer process, and $S_c$ their electron-phonon coupling strength. The free energy change upon electron transfer, $\Delta G_{PeT}$ is given by:

$$\Delta G_{PeT} = \Delta G_0 - \Delta\vec{\mu}.\vec{E} \tag{S8}$$

where $\Delta\vec{\mu}$ is the change in the electric dipole moment of the donor-acceptor pair when electron transfer occurs. The reverse reaction rate constant is simply given by:

$$k_{21} = k_{12}\exp\left(\frac{\Delta G_{PeT}}{k_B T}\right) \tag{S9}$$

and is negligible if $\Delta G_0 < 0$. The important ingredient of this model is that the free energy change $\Delta G_{PeT}$ depends linearly on the transmembrane electric field $\vec{E}$ (Eq. (S8)).

Rates $k_{21}$ ($\ll k_{12}$) and $k_{20}$ are electric field-dependent while $k_{10}$ is (in general) unaffected by it. A change in MP therefore induces a change in $\Delta G_{PeT}$ proportional to the change in the local electric field, which results into a change in the PeT kinetic rate $k_{12}$.(11,12) As shown by Li, both lifetime components of the decay depend in a non-trivial manner on the electric field.(10) However, when $k_{21} \ll k_{20}$, the fluorescence quantum yield $Y_f$ is given by the simple expression:

$$Y_f \sim \frac{k_f}{k_{10}}\left(1+\frac{k_{12}}{k_{10}}\right)^{-1} \tag{S10}$$

where $k_f$ is the radiative rate of de-excitation from 1 to 0. The amplitude-averaged lifetime is then:

$$\langle \tau \rangle_a = \frac{Y_f}{k_f} \sim k_{10}^{-1}\left(1+\frac{k_{12}}{k_{10}}\right)^{-1} \tag{S11}$$

This expression therefore indirectly provides the electric field dependency of the measured amplitude-averaged lifetime.

#### Supplementary Note 7: Fluorescence intensity profile analysis for stained bacterial membrane

If the membrane can be described as a cylinder of radius $R$ with the $y$ axis as its axis of symmetry (Supplementary Figure 16) and the PSF can be approximated by a 3D Gaussian with parameters $\sigma_{xy}$ and $\sigma_z$, it is straightforward to verify that the collected intensity at position ($x_0$, 0, 0) is proportional to:

$$I(x_0,0,0)=2\pi\sigma_{xy}\sigma_*\exp\left(-\frac{R^2}{2\sigma_z^2}\right)\exp\left(\frac{x_*^2}{2\kappa^2\sigma_*^2}\right)\left(erf\left(\frac{x_*+R}{\sqrt{2}\sigma_*}\right)-erf\left(\frac{x_*-R}{\sqrt{2}\sigma_*}\right)\right) \tag{S12}$$

where:

$$x_*=-x_0(1-\kappa^{-2})^{-1}$$
$$\sigma_*=\frac{\sigma_{xy}}{\sqrt{1-\kappa^{-2}}} \tag{S13}$$
$$\kappa=\sigma_z/\sigma_{xy}$$

κ can also be estimated using the ratio between lateral and axial resolution given by the Rayleigh criterion in a confocal microscope (e.g. (13)):

$$\delta_{xy}=0.51\frac{\lambda}{NA}$$
$$\delta z=0.88\frac{\lambda}{n-\sqrt{n^2-NA^2}}, \tag{S14}$$

where $\lambda$ is the excitation wavelength, $n$ the index of refraction of the immersion medium and NA the objective lens numerical aperture. It follows that:

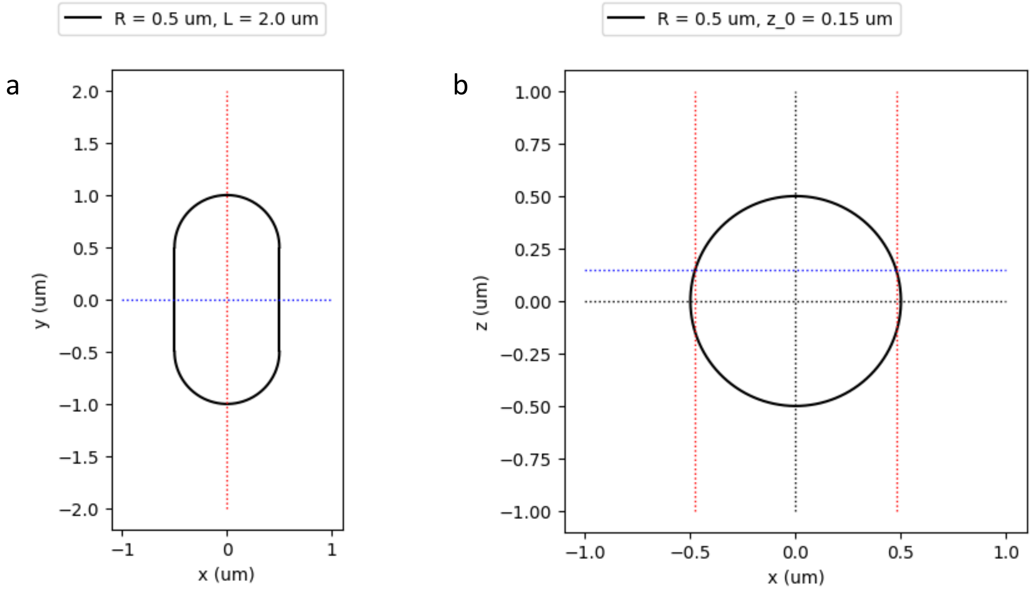

**Supplementary Figure 16**: Bacterial intensity profile geometric definitions. The bacterium is assumed to be comprised of a cylindrical body of length $L-2R$, with two hemispherical caps. (a) Top view, (b) front view. The intensity profile is computed along the $x$ direction (blue dotted line in a and b) in the middle of the cell ($y=0$) in order for the influence of the caps to be negligible. The cross-section can be obtained at a $z$ depth (blue dotted line in b) different than the mid-plane (indicated by a horizontal black dotted line in b).

$$\kappa=\frac{\delta_z}{\delta_{xy}}=\frac{1.73}{\alpha-\sqrt{1-\alpha^2}} \tag{S15}$$
$$\alpha=NA/n$$

$\sigma_{xy}$ can be taken as the parameter of the Gaussian approximation to the Airy disk:

$$\sigma_{xy} = 0.21\frac{\lambda}{NA} \tag{S16}$$

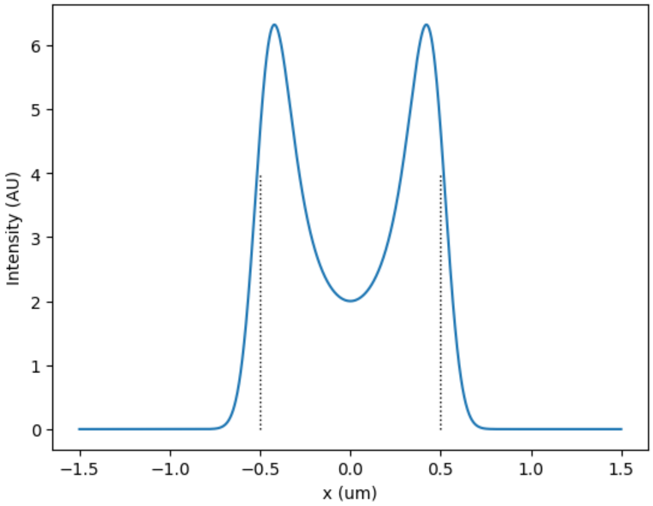

**Supplementary Figure 17**: Intensity profile at the midplane computed using the Google Colab notebook bacterial-membrane-intensity-profile.ipynb with the parameters mentioned in the text. (1)

For instance, for $\lambda$ = 514 nm, NA = 1.4 and $n$ = 1.52, we obtain $\sigma_{xy}$ = 77 nm, $\sigma_z$ = 251 nm, $\alpha$ = 0.92, $\kappa$ = 3.25. The corresponding profile for $R$ = 0.5 µm is represented in Supplementary Figure 17, and shows that the intensity in the middle of the cell (where no dye is present), is nonzero due to the signal collected from the membrane below and above the focal plane.

The intensity profile at a depth $z_0$ other than the midplane cannot be expressed in as simply but can be calculated numerically according to:

$$I(x_0,0,z_0) = \sqrt{2\pi}\sigma_{xy}R\exp\left(-\frac{\left(R^2+z_0^2\right)}{2\sigma_z^2}\right)\exp\left(\frac{x_*^2}{2\kappa^2\sigma_*^2}\right)\int_0^\pi d\theta\sin\theta\exp\left(-\frac{\left(R\cos\theta+x_*\right)^2}{2\sigma_*^2}\right)2\cosh\left(\frac{z_0R\sin\theta}{\sigma_z^2}\right) \tag{S17}$$

An example the effect of such a defocusing on the contrast between the brighter regions of the membrane and the inside of the membrane is shown in Supplementary Figure 18. In particular, the apparent width of the cell (distance

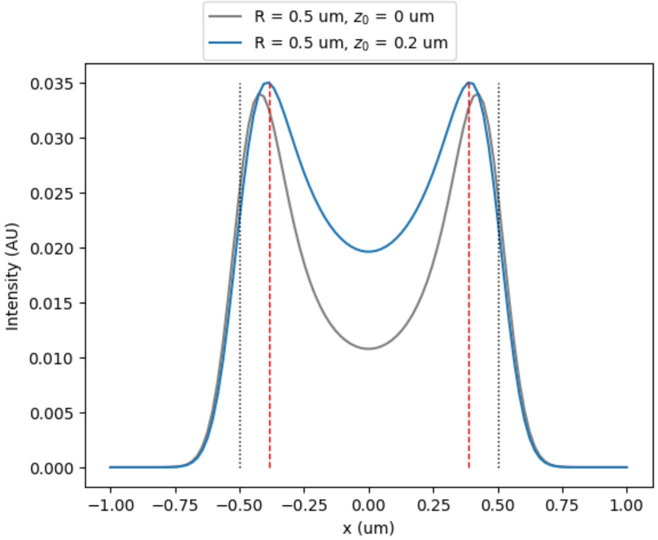

**Supplementary Figure 18**: Effect of defocusing on the intensity profile. The gray curve corresponds to the profile acquired at the mid-plane (identical to that in Suppl. Figure 9), which the blue curve corresponds to the profile acquired with a defocusing of 0.2 µm. The parameters used for the calculation (reproducible using he Google Colab notebook bacterial-membrane-intensity-profile.ipynb) (1) are mentioned in the text. The dotted gray vertical lines indicate the location of the membrane at the mid-plane ($x = \pm 0.5$ µm), while the dashed red lines indicate the location of the intensity maxima ($x = \pm 0.39$ µm).

separating the intensity maxima, indicated by red dashed vertical lines in the figure) decreases as focus is lost, while the intensity increases slightly, before decreasing as the amount of defocus increases.

Examples of the use of this model with real data are presented in Figure 2 in the main text, an extended version of which is shown in Supplementary Figure 19.

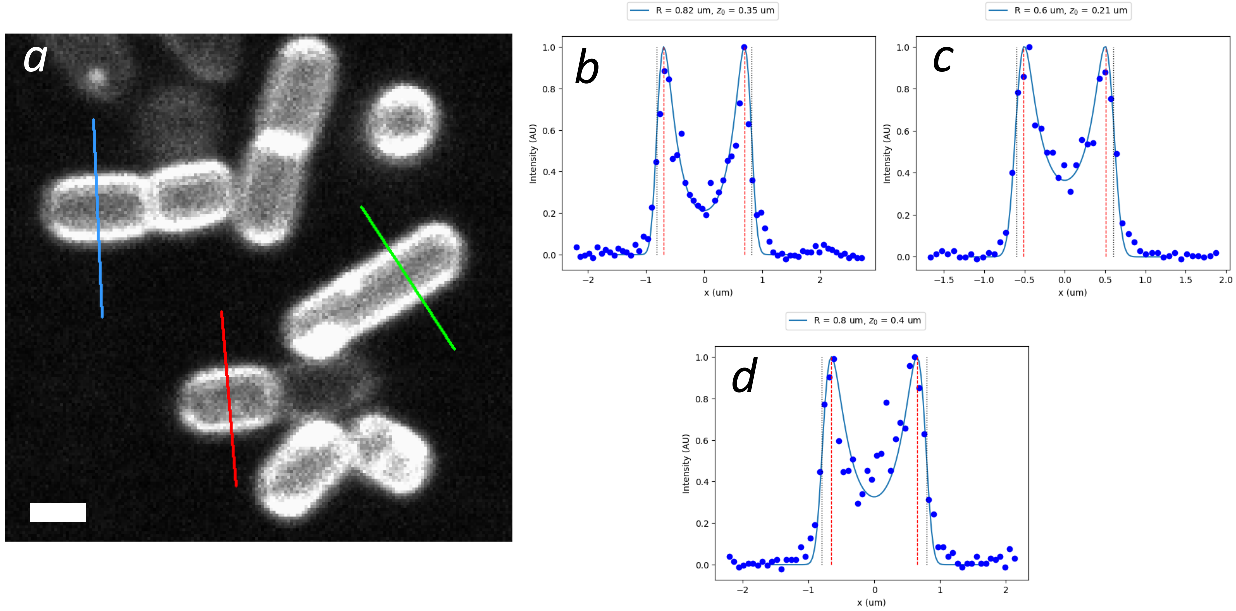

**Supplementary Figure 19**: Experimental intensity profiles. (a) *B. subtilis* cells were grown in resuspended in M9 media and stained with VF2.1.Cl and imaged using a FLIM confocal setup as described (pixel size: 72 nm/pixel, scale bar: 1 µm). Intensity profiles along the 3 segments are shown in b (blue), c (green) and d (red) as blue dots. The blue curves correspond to Eq. (S17) with parameters $R$ and $z_0$ adjusted for best fit indicated in the legend. It is noteworthy that the cells appear to be located at different focal planes ($z_0$ = 0.35, 0.21, 0.4 µm) and characterized by different radii ($R$ = 0.82, 0.6, 0.8 µm). It is possible that additional degrees of freedom (e.g. tilt angle) might need to be considered to better account for the observed diversity of bacterial shape parameters.

#### Supplementary Note 8: Estimation of the potential drop across a semipermeable membrane in the presence of a membrane potential

In this note, we compute the relation between potassium Nernst potential (from which the membrane potential can be determined) and the electric field in the membrane, to which the VF probe is exposed and which directly affects the probe's fluorescence lifetime. Note that we completely neglect the contribution of the bacterial cell wall (~34 nm in *B. subtilis*), which most likely harbors a gradient of ions across its thickness and therefore contributes an additional drop or increase of electric potential between the bulk of the solution and the membrane. (14)

The electrostatic potential across a semi-permeable planar membrane separating electrolyte solutions has been extensively studied in the literature. We follow the simplified analysis presented in ref. (15), with the difference that the two sides of the membrane contain different electrolytes. The applied potential difference $V$ in ref. (15) is assumed to be equal to the membrane potential $V_m$ obtained from the Donnan or GHK solution, as discussed in the text.

Noting all "outside" quantities with index *out*, "inside" quantities with index *in* and membrane-related quantities with index *m*, we have the following solutions for the electrostatic potential $\phi(z)$ (axis $z$ is oriented from outside the cell to inside the cell, with the membrane boundaries located at $z = 0$ and $z = d$):

$$\begin{cases} \phi_{out}(z) = A_{out} e^{\kappa_{out} z} \\ \phi_m(z) = A_m + B_m z \\ \phi_{in}(z) = V - A_{in} e^{-\kappa_{in} z} \end{cases}, \tag{S18}$$

with the following continuity relations for the potential:

$$\begin{cases} \phi_{out}(-\infty) = 0 \\ \phi_{in}(+\infty) = V \\ \phi_{in}(0) = \phi_m(0) \\ \phi_m(d) = \phi_{in}(d) \end{cases}. \tag{S19}$$

and for the electric field:

$$\begin{cases} \varepsilon_W \phi_{out}'(0) = \varepsilon_m \phi_m'(0) \\ \varepsilon_W \phi_{in}'(d) = \varepsilon_m \phi_m'(d) \end{cases}, \tag{S20}$$

where $\varepsilon_W, \varepsilon_m$ are the relative electric permittivity of water and the membrane respectively, and $\kappa_i$ the Debye-Hückel factor of medium *i*.

The solution is given by:

$$\begin{cases} A_{out} = A_m = \left(1 + \dfrac{\kappa_{out}}{\kappa_{in}} + \dfrac{\varepsilon_W}{\varepsilon_m}\kappa_{out} d\right)^{-1} \\ B_m = \dfrac{\varepsilon_W}{\varepsilon_m}\kappa_{out} A_{out} \\ A_{in} = \dfrac{\kappa_{out}}{\kappa_{in}} e^{\kappa_{out} d} A_{out} \end{cases}. \tag{S21}$$

The potential drop across the membrane, $V_{mb} = \phi_m(d) - \phi_m(0)$ is given by:

$$V_{mb} = \frac{\varepsilon_W}{\varepsilon_m}\kappa_{out} d \left(1 + \frac{\kappa_{out}}{\kappa_{in}} + \frac{\varepsilon_W}{\varepsilon_m}\kappa_{out} d\right)^{-1} V. \tag{S22}$$

Introducing the notations:

$$\begin{cases} \alpha = \dfrac{1 + \dfrac{\kappa_{out}}{\kappa_{in}}}{1 + \dfrac{\kappa_{out}}{\kappa_{in}} + \gamma} \\ \gamma = \dfrac{\varepsilon_W}{\varepsilon_m}\kappa_{out} d \end{cases}, \tag{S23}$$

the difference between $V$ (set by the Donnan or GHK solution) and $V_{mb}$ (which sets the amplitude of the electric field within the membrane as $E = V_{mb}/d$) is equal to:

$$V - V_{mb} = \alpha V \ . \tag{S24}$$

To obtain an order of magnitude of this difference, we need to estimate the Debye-Hückel factors $\kappa_i$ :

$$\kappa_i^2 = \frac{e^2}{k_B T \varepsilon_0 \varepsilon_i} \sum_j z_j^2 C_j = \frac{2e^2}{k_B T \varepsilon_0 \varepsilon_i} I_i \tag{S25}$$

where $C_j$ is the concentration of ion $j$ (in medium $i$) and $z_j$ its charge in unit of electric charge. $\kappa_i^2$ is proportional to the ionic strength $I_i$, allowing to rewrite $\alpha$ as:

$$\begin{cases} \alpha == \dfrac{1+\beta}{1+\beta+\gamma} \\ \beta = \sqrt{I_{out}/I_{in}} \end{cases} \tag{S26}$$

Using the composition of MSgg given above for the outside electrolyte, we obtain $I_{out}$ = 65 mM. Assuming that the internal ionic strength is dominated by $K^+$, we can set $I_{in}$ = 300 mM. It follows that $\kappa_{out}^{-1} = 12\,\mathring{A}$ and $\kappa_{in}^{-1} = 5.6\,\mathring{A}$. Using $\varepsilon_W$ = 80.2 and $\varepsilon_m$ = 6 (16,17), we obtain $\gamma$ = 55.5 , $\alpha$ = 0.026, indicating a small but measurable difference between the calculated Nernst potential and the potential drop across the membrane. The difference is reduced to ~1.6% in the case of identical ionic strength on both sides of the membrane. The corresponding electric potential profile is plotted in Supplementary Figure 20.

Note that the concentration of the different species $j$ , $C_j^{(i)}$ on each side ($i$ = 1 or 2), can be computed according to:(15)

$$\begin{cases} C_j^{(1)}(z) = C_j^{(1)}(-\infty)\left(1 - \dfrac{z_j e}{k_B T}\phi_{out}(z)\right) \\ C_j^{(2)}(z) = C_j^{(2)}(+\infty)\left(1 - \dfrac{z_j e}{k_B T}\phi_{in}(z)\right) \end{cases} \tag{S27}$$

These equations indicate that the concentrations differ from the bulk concentrations only within a narrow layer of typical dimension given by the Debye length of the corresponding medium.

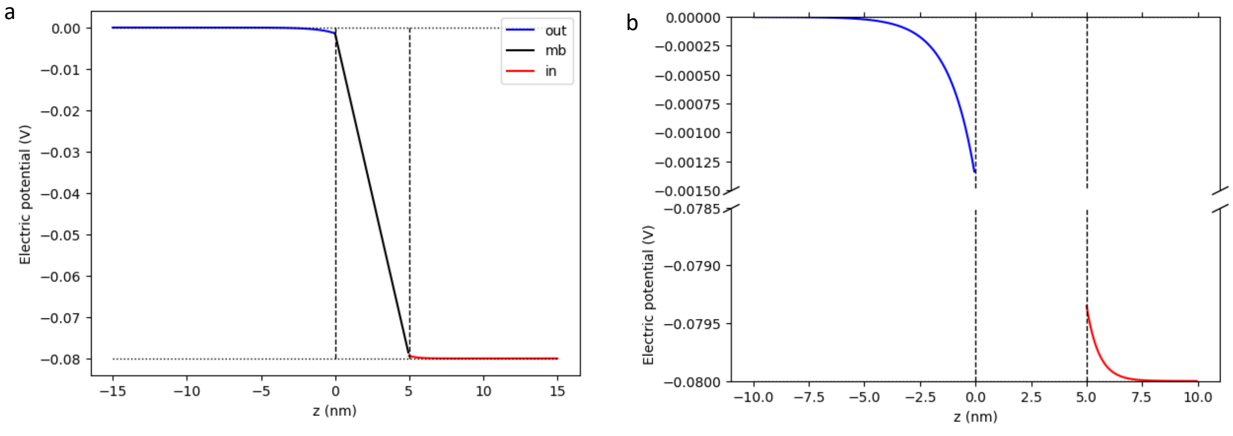

**Supplementary Figure 20**: Electrostatic potential across the membrane according to the model discussed here. Model parameters: $I_{out}$ = 65 mM, $I_{in}$ = 65 mM, $d$ = 5 nm. (a) The outside potential drops from 0 to -1 mV within a few nm of the membrane (blue curve), followed by a linear decrease close to the total potential drop (-0.8 V) within the membrane (black segment), followed by a small and rapid final drop in the vicinity of the membrane inside the cell (red curve). (b) Details of (a) in the vicinity of the membrane boundaries. The curves were computed using the Google Colab notebook membrane-potential.ipynb.(2)

#### Supplementary Note 9: Additional calibration analysis details

The $V_m$ *vs* <τ> calibration curve of Figure 4c relies on heuristical fits of two sets of experimental data: 1) the reported $V_m$ *vs* $[K^+]_{out}$ data of Shioi *et al.* obtained using radioactive Nernstian probes, and 2) the <τ> *vs* $[K^+]_{out}$ obtained in this work. We did not try to fit the data to any *a priori* theoretical model, but tried instead to obtain an analytical formula that would allow a simple conversion of <τ> into $V_m$, over the range of <τ> obtained in our experiments.

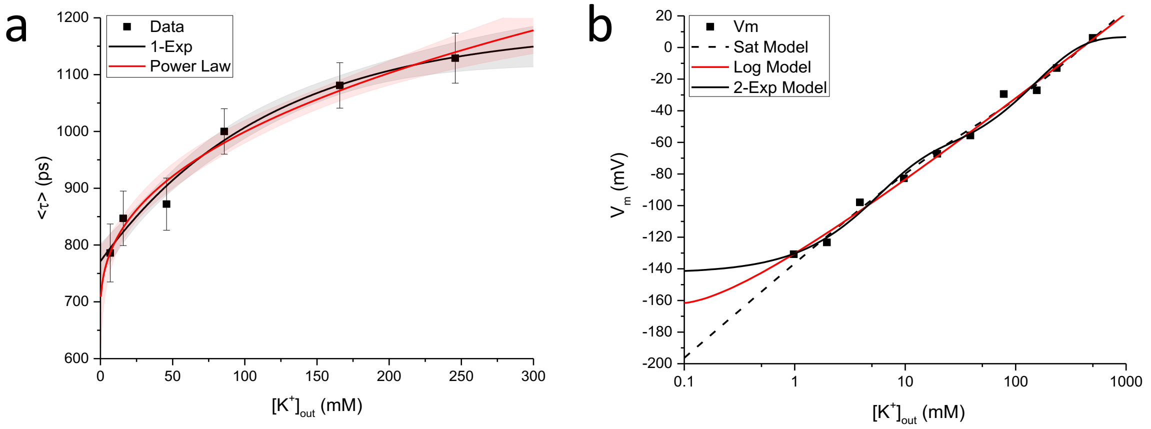

**Supplementary Figure 21**: Heuristic models used to fit our calibration series (a) and Shioi *et al.*'s calibration data (b). The different models are discussed in the text of this supplementary note.

To model our data (Figure 4b), we tested two simple analytical models: (i) a single-exponential saturation model, and (ii) a power-law model with offset. Both are represented in Supplementary Figure 21a. and satisfactorily account for the data, with one major difference: the exponential model only outputs lifetimes in the [771, 1149] ps range, while the power law model outputs lifetimes in the [678,+∝[ range. We therefore used the power law model, given by:

$$\langle\tau\rangle = F_1\left(\left[K^+\right]_{out}\right) = \langle\tau\rangle_0 + A\left[K^+\right]_{out}^{\alpha} \quad \text{(S28)}$$

with $<\tau>_0$ = 678.28 ps, $A$ = 50.26, $\alpha$ = 0.4027. This relation can be easily inverted as:

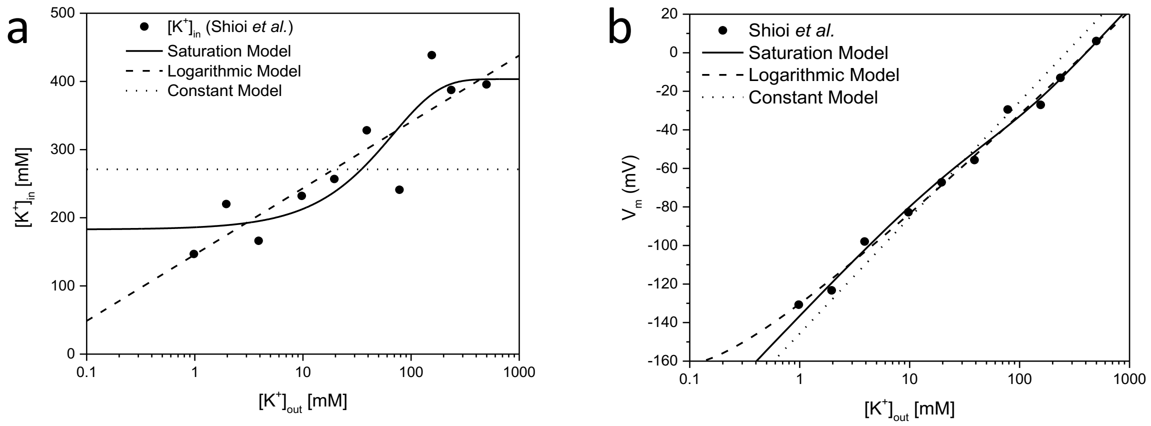

**Supplementary Figure 22**: *B. subtilis* membrane potential as function of external $K^+$ concentration according to Shioi *et al.*.(3) (a) $[K^+]_{in}$ vs $[K^+]_{out}$ as measured by Shioi *et al.* $[K^+]_{in}$ values were obtained from Figure 3 in ref. (3), which represents $V_m$ as a function of $[K^+]_{out}$, using the Nernst potential equation. The plain curve is the best fit to the data of a saturation model (Eq. (S30) ). The dashed curve is a fit to a logarithmic model, while the dotted curve is a fit to a constant. (b) $V_m$ measured with radioactive Nernstian probes as a function of external $K^+$ concentration $[K^+]_{out}$, adapted from Figure 3 in ref. (3). The curves correspond to the heuristic models shown in (a).

$$\left[K^+\right]_{out}\left(\langle\tau\rangle\right) = F_1^{-1}\left(\langle\tau\rangle\right) = \left[\frac{\left(\langle\tau\rangle - \langle\tau\rangle_0\right)}{A}\right]^{-1/\alpha} \quad (S29)$$

To model Shioi *et al.*'s data, we tried 3 different models: (i) a saturation model for the internal potassium concentration $[K^+]_{in}$ *vs* $[K^+]_{out}$, (ii) a logarithmic model for $[K^+]_{in}$ *vs* $[K^+]_{out}$ and (iii) a bi-exponential model for $V_m$ *vs* $[K^+]_{out}$. The saturation model stems from observing the relation between internal and external potassium concentrations derived from Shioi *et al.*'s $V_m$ *vs* $[K^+]_{out}$ data (derived from Figure 3 in ref. (3)), shown in Supplementary Figure 22a. This relation can be fitted by the following heuristic law:

$$\left[K^+\right]_{in} = \left[K^+\right]_{in}^{\infty} + k\exp\left(-\left[K^+\right]_{out} / \left[K^+\right]_{out}^{*}\right) \quad (S30)$$

where $[K^+]_{in}^{\infty} = 403.3$ mM, $k = -220.6$ mM and $[K^+]_{out}^{*} = 68.0$ mM. This provides a better fit to their $V_m$ *vs* $[K^+]_{out}$ data (Supplementary Figure 22b, plain curve) than the assumption made by the authors of a constant internal potassium concentration (best fit: $[K^+]_{in} = 271$ mM, Supplementary Figure 22b, dotted curve). A simpler logarithmic relation between $[K^+]_{in}$ *vs* $[K^+]_{out}$ is also shown as a dashed curve.

The resulting $V_m$ *vs* $[K^+]_{out}$ curves are represented in Supplementary Figure 21b. The saturation model, represented as a dashed curve, has the inconvenience of not bounding the range of membrane potential values at low or high external potassium concentrations, potentially resulting in unphysical extrapolated values when combined with the $<\tau>$ *vs* $[K^+]_{out}$ data. The logarithmic model (represented as a red curve in Supplementary Figure 21b) doesn't suffer from unbounded lower values, but is not bounded as far as high $V_m$ values are concerned. For all these reasons, we decided to fit the $V_m$ *vs* $[K^+]_{out}$ data of Shioi *et al.* by a bi-exponential model (black curve in Supplementary Figure 21b) using the following form and parameters:

$$V_m\left(\left[K^+\right]_{out}\right) = F_2\left(\left[K^+\right]_{out}\right) = V_0 + A_1\exp\left(-\left[K^+\right]_{out} / K_1\right) + A_2\exp\left(-\left[K^+\right]_{out} / K_2\right) \quad (S31)$$

with $V_0 = 6.71$, $A_1 = -70.41$, $K_1 = 5.37$, $A_2 = -79.0$, $K_2 = 151.42$.

In summary, the heuristic calibration form used in this work is:

$$V_m\left(\langle\tau\rangle\right) = F_2\left(F_1^{-1}\left(\langle\tau\rangle\right)\right) \quad (S32)$$

where $F_1^{-1}$ and $F_2$ are defined by Eq. (S29) & (S31) respectively. This relation is represented in Supplementary Figure 23a (reproduced in Figure 4c in the main text). It is visually well approximated by two piecewise linear functions, with a slope of 0.2 mV/ps (5 ps/mV) in the [850 ps – 1.2 ns] range, and increasing to up to 0.8 mV/ps (1.2 ps/mV) for lower lifetimes (lower external $K^+$ concentration or higher membrane hyperpolarization). This is confirmed by looking at the derivative of $V_m(<\tau>)$, represented in Supplementary Figure 23b, in which the two local slopes are indicated by dashed lines.

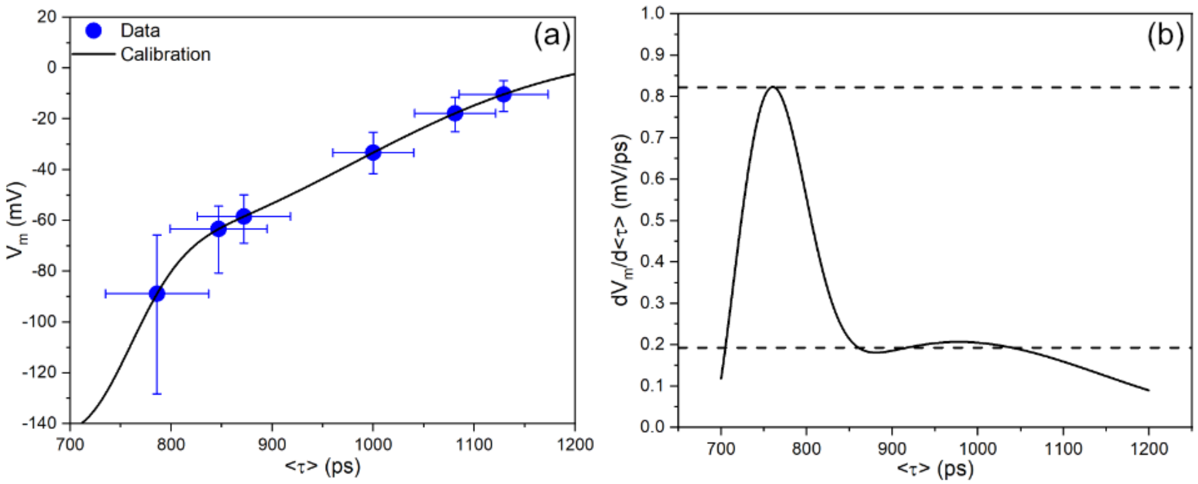

**Supplementary Figure 23**: (a): $V_m(<\tau>)$ calibration curve and (b): its derivative, with the two slopes of the piecewise linear approximation are indicated by dashed lines.

#### Supplementary Note 10: Fluorescence lifetime measurements outside *B. subtilis* cells

Because the signal-to-background ratio in the current measurement was fairly low (e.g. SBR = 3.7 in the valinomycin MSgg data used in the calibration series of Figure 5), we investigated the characteristics of the fluorescence decay measured in the "background" areas. The existence of a background fluorescence signal can be due to a variety of sources, among which the presence of free dye in solution (the dye being free to leave the membrane) or freely moving bacteria coming and going out-of-focus during data acquisition are the most likely.
The average lifetime of VF2.1.Cl in different conditions is discussed in the text and reported in Supplementary Figure 3 and Supplementary Table 3.
The average lifetime of out-of-focus cells should be identical to that measured in focus (and lower than that of the free dye).
In summary, we expect the background fluorescence to be a linear combination of these two potential sources.
To characterize the background fluorescence decays, we first defined background ROIs by first merging all cell ROIs used in the cell MP analysis, adding "bright" or "hazy" ROIs suspected to correspond to out-of-focus cells or cell clusters, and then taking the complementary ROI of all these regions. The phasor of this background ROI was computed for one FOV per condition studied in the calibration series and the corresponding amplitude-averaged lifetime computed using the same references used for the cell fluorescence analysis.
Supplementary Figure 24 shows the observed behavior of the background fluorescence lifetime (blue triangles), together with the corresponding mean and standard deviation of the cell lifetimes (black squares). Both are somewhat correlated, the background lifetime being larger than that observed in cells, but lower than that of the free dye (as expected).

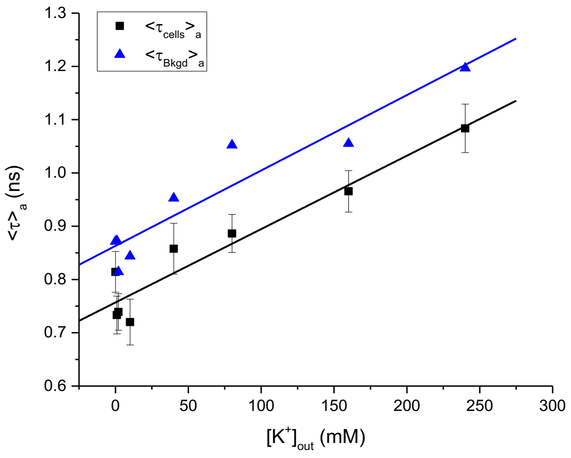

**Supplementary Figure 24**: Comparison of amplitude-averaged lifetime of the background fluorescence (blue triangles) and of VF2.1.Cl in *B. subtilis* (black square) in MSgg + 25 µM valinomycin + KCl at different concentrations. The lines are linear fits to the data.

Focusing on the last dataset (cells in MSgg + 25 µM valinomycin + 240 mM KCl), Supplementary Figure 25 shows the corresponding phasor plot with the background fluorescence phasor (black circle, corresponding to a single ROI) and cell phasors (blue dots) as well as the phasor of the pure dye in the same MSgg + 25 µM valinomycin + 240 mM KCl conditions (teal square). Supplementary Figure 25a also represents the phasor references used for amplitude-averaged lifetime analysis (green and red circles, localized at the intersection of the green and red vertical and horizontal lines, respectively). The background phasor (black circle) is clearly distinct from the majority of *B. subtilis* cell phasors (blue dots) and located closer to the long lifetime reference (red dot) than the cell phasors are, consistent with a larger average lifetime. The open red and green squares located on the UC correspond to the single exponential components of the pure dye decay, and are close to the single-exponential phasor references used for the *B. subtilis* cell signal analysis (green and red lines intersections).

Supplementary Figure 25b focuses on the background phasor (black circle) and compares it to the pure dye phasor (teal square, where the red lines now intersect) and the average of the *B. subtilis* cell phasors (pink open square, where the green lines now intersect). All these phasors appear aligned along the segment connecting the single-exponential components of the pure dye decay (only the longer lifetime component is visible in b as a red square), suggesting indeed that the background phasor ($z_{Bkgd}$) can be interpreted as a linear combination of the phasor of the pure dye in solution ($z_{Dye}$) and the average phasor of the dye in *B. subtilis*' membrane ($z_{Mb}$):

$$z_{Bkgd} = 0.78 z_{Mb} + 0.22 z_{Dye} \quad \text{(S33)}$$

Measurements in the absence of mobile cells in suspension and in the presence of constant flow of dye-free medium would therefore eliminate this contribution and increase the SBR.

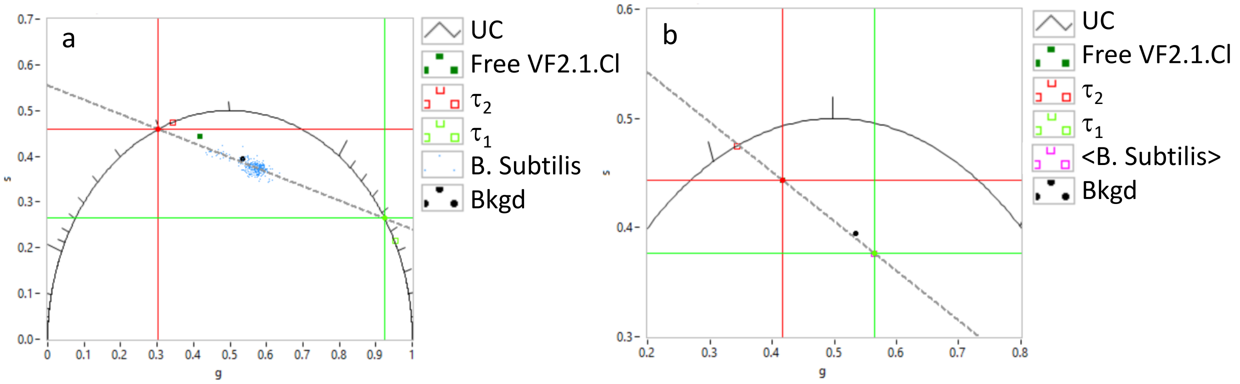

**Supplementary Figure 25**: Phasor analysis of background fluorescence. (a) Phasor of *B. subtilis* cells (blue scatterplot), background (black circle) and free dye (teal square). The phasor references used for the analysis of the *B. subtilis* cells and background are indicated by the intersection of the red and green lines with the UC. The green and red open squares on the UC correspond to the single-exponential components of the free dye's decay. (b) Zoom on the background phasor (black circle), average of *B. subtilis* phasors (pink open square/green cursor) and free dye (teal square/red cursor). The background phasor can be interpreted as a linear combination of the cells' average phasor and the free dye's phasor.

#### Supplementary Note 11: Literature reporting estimated membrane potential values for B. subtilis

| Literature | Strain, Genotype | Extracellular medium | Membrane Potential (mV) | Method of measurement |
|---|---|---|---|---|
| Hosoi et al., Biochim Biophys Acta 1980, 600 (3), 844-852 | BC26, pheA12 argA3 ery | Basal solution (a minimal media with 0.5% Glycerol, 0.1 mM EDTA), pH 7.0 | -120 to -80 | Direct, Nernst equilibrium for [TPMP+], microelectrode-based measurement of partitioning of [TPMP+] (a radioactive membrane permeable cation) across the membrane |
| Zaritsky et al., J Membr Biol 1981, 63 (3), 215-231 | OI1, ilv C1 leu-1 | Modified chemotaxis media (a minimal media with 0.5% Glycerol, 0.1 mM EDTA), pH 7.5 | -120 | (i) Direct, measurement based on Nernst equilibrium for [TPMP+] and it's partitioning of [TPMP+] across the membrane; (ii) Indirect, measurement based on partitioning of DiSC3(5) and it's relative fluorescence intensity change. Valinomycin together with excess [K+] used to modulate [K+] electrochemical gradient/K+ Nernst equilibrium. |
| Shioi et al., J Bacteriol 1980, 144 (3), 891-897 | BC26, pheA12 argA3 ery | Motility media (a minimal media 10 mM potassium succinate, 0.1 mM EDTA), pH 7.5 | -140 @ pH 7.4<br>-125 @ pH 7.0<br>0 @ pH 4.5 | Direct, measurement based on Nernst equilibrium for [TPMP+] and it's partitioning of [TPMP+] across the membrane. Valinomycin together with excess [K+] used to modulate [K+] electrochemical gradient/K+ Nernst equilibrium. Membrane potential estimated based on Nernst equilibrium for [TPMP+] matches quite well theoretically calculated membrane potential based on Nernst equilibrium for [K+] (intracellular [K+] estimated to be 450 mM) |
| Winkel et al., Front Cell Dev Biol 2016, 4, 29 | 168, trpC2 | Chemically defined media (a minimal media with 2.2 mM D-Glucose, 2.1 mM L-Glutamic acid) | -110 | Indirect, measurement based on partitioning of DiSC3(5) and it's relative fluorescence intensity change. Valinomycin together with excess [K+] used to modulate [K+] electrochemical gradient/K+ Nernst equilibrium. Membrane potential calculated based on Nernst equilibrium for [K+] (intracellular [K+] taken as to be 300 mM) |
| Present study | 168, trpC2 | MSgg (a media with 68.4 mM (0.5% v/v) Glycerol, 29.5 mM (0.5% wt/v) Glutamate), pH 7.0<br>M9 (a minimal media with 0.4% wt/v Glucose, pH 7.0 | -65<br>-127 | Indirect, measurement based on fluorescence lifetime change of membrane embedded VF2.1.Cl. Valinomycin together with excess [K+] used to modulate [K+] electrochemical gradient/K+ Nernst equilibrium. Membrane potential calculated based on Nernst equilibrium for [K+] (intracellular [K+] taken as to be 300 mM) |

#### Supplementary Note 12: 'Puncta' in membrane labelling

The 'puncta' in membrane labelling i.e. punctuated/irregular fluorescence intensity patterns are not caused by staining with the VoltageFluors. During sample preparation we first stained bacterial cells with VF2.1.Cl and subsequently divided the cells into three groups: (a) unperturbed (*i.e.* resuspended in MSgg media only), (b) perturbed with 25 μM valinomycin alone and (c) perturbed with 25 μM valinomycin + 240 mM KCl (excess KCl).
The formation of distinct punctuated patterns of fluorescence intensity were observed after exposing the cells to chemically depolarizing conditions (b) and (c), but not in unperturbed condition (a) (see Supplementary Figure 26). They were only visible under confocal fluorescence imaging of VF2.1.Cl, but not in white light imaging (Supplementary Figure 27). This suggests that puncta are membrane (or membrane-bound) features and not sub-cellular substructures such as vesicles, which should be visible under white light illumination, if larger or comparable in size with the diffraction limit of the objective lens used (~150 nm). As just mentioned, punctuated patterns of fluorescence intensity were not observed when the cells were resuspended in MSgg only (unperturbed, condition (a), Supplementary Figure 26a). In other words, the formation of these punctuated patterns is not associated with membrane labeling, but with perturbation (with valinomycin in our experiments), and appears exacerbated by stressful conditions such as excess external potassium.

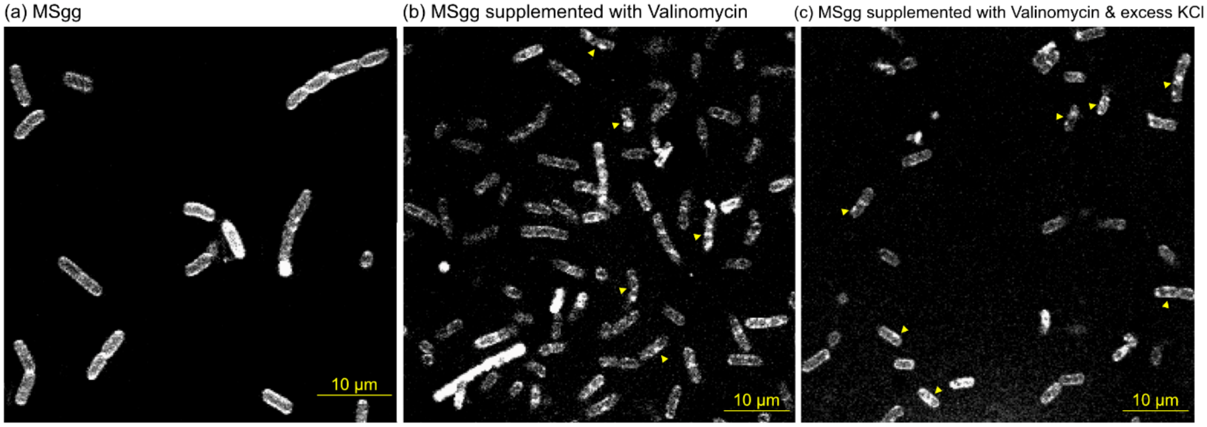

**Supplementary Figure 26**: Fluorescence intensity images of B. subtilis cells stained with VF2.1.Cl at different extracellular chemical conditions: distinct punctuated fluorescence intensity patterns were observed when cells are perturbed with valinomycin and KCl (b)-(c). Some cells of interest are indicated with yellow marker to point at examples of puncta. Similar patterns were not observed when cells are unperturbed (a).

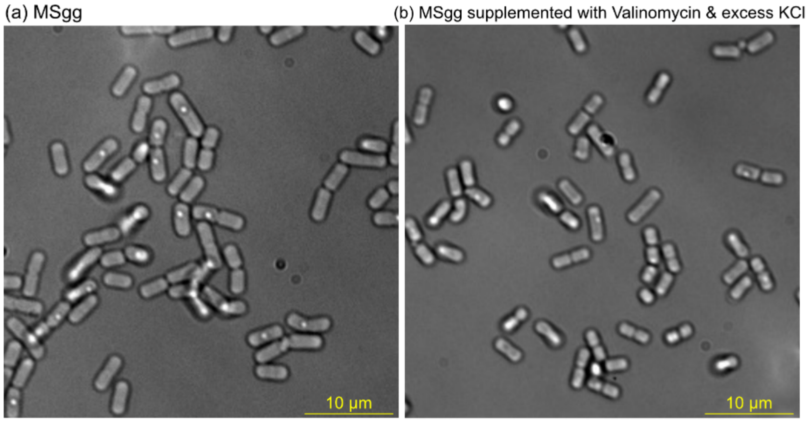

**Supplementary Figure 27**: White light images of B. subtilis cells in different extracellular chemical conditions: no specific cellular substructures (apart from spores, some of which are visible in these images, and are not stained by VF2.1.Cl) similar to punctuated fluorescence intensity patterns, have been observed irrespective of extracellular chemical conditions.

Puncta have been observed in bacterial membranes in other circumstances, unrelated to membrane labeling but revealed by it. They have been attributed to the presence of membrane patches/microdomains with different local properties such as membrane fluidity. For instance, it has been shown that there exists membrane microdomains/regions of increase fluidity (RIFs) in rod-shaped Gram-positive and Gram-negative bacteria (18-20). These microdomains likely contain clusters of lipids in a fluid disordered state. When wild-type B. subtilis cells were stained with DiI-C12, a lipophilic dye that selectively localizes within lipid membranes, a distinct punctuated fluorescent pattern characteristic of RIFs was observed. These RIFs act as platform to organize various proteins of importance (such as peripheral membrane proteins (PlsX), cell division proteins (MinD, FtsA), bacterial actin homologue (MbI, MreB, MreBH) for crucial physiological processes. Interestingly, the localization pattern of proteins like MreB (18) and PlsX (20) in B. subtilis (unperturbed), was observed to overlap with the punctuated distribution of DiI-C12. However, in B. subtilis mutant strains lacking mreB, mbl, and mreBH, DiI-C12 labels membrane uniformly without any patches. This indicates that RIFs are not observed or formed in these strains (18).

Others have argued that the shape of proteins or protein complexes within the membrane can influence the local curvature, leading to a disruption of diffusion and resulting in a patterned (or punctuated) distribution of membrane proteins (21). It is important to note that not all membrane proteins localize in a punctuated pattern in an uninhibited bacterial cell. Often, proteins exhibit a punctuated pattern when subjected to some physiological stress, such as artificial modulation of membrane potential. Strahl and Hamoen's research (18,22) has revealed that modulation of membrane potential significantly affects the spatial organization of several key cytoskeletal and cell division proteins, including MinD, FtsA, and MreB. Disruption of membrane potential using ionophores (CCCP, valinomycin), antibiotics (nisin), or bacteriocins (colicillin N) triggers abnormal localization of these proteins. MinD loses its polar localization, FtsA loses its septal localization, and MreB loses its helical distribution around the cell. Interestingly, in the case of MinD and MreB, a punctuated fluorescent pattern is observed.

In a separate study, Strahl and Hamoen have observed that when B. subtilis stained with Nile Red is subjected to membrane potential-dissipating conditions, its cell membrane undergoes a transformation from uniform labeling to a non-uniform, punctuated fluorescent pattern. Nile Red, a membrane dye, partitions into lipid membranes based on its intrinsic hydrophobicity, similar to our voltage probe VF2.1.Cl. Nile Red's fluorescence is sensitive to physical changes in the lipid environment.

Therefore, a punctuated membrane staining of Nile Red simply indicates inhomogeneities in the membrane's lipid environment due to membrane potential-dissipating conditions. When B. subtilis cells stained with a fluidity-sensitive dye, Laurdan, membrane potential dissipation was observed to be associated with the formation of membrane microdomains with increased membrane fluidity. These microdomains were colocalized with the punctuated Nile Red stain.

During analysis we have treated each individual cell as a single entity, obtaining a single lifetime from each cell, irrespective of the presence or absence of punctuated/irregular fluorescence intensity patterns or other possible morphological features. As discussed above, puncta are commonly observed in physiologically stressed cells, and would have therefore been present in the calibration measurements performed by Shioi *et al.* (and in other calibration measurements) in which they used a Nernstian probe to measure the membrane potential. In this respect, our analysis is already much more fine-grained than theirs, since we obtain a distribution of lifetimes in each condition, compared to a single value in the work of Shioi *et al.* Our measured lifetime distributions are well-described by Gaussian (i.e. normal) distributions, which allows us to use the mean observed lifetime for correlation with the Shioi *et al.* work.
